# Dual inhibition of the nonsense mediated mRNA decay enhances tumour immunogenicity, drives immunoediting, and potentiates checkpoint blockade

**DOI:** 10.64898/2025.12.04.692418

**Authors:** Ivan Zadra, Davor Orsolic, Dimitri Kounis, Mario Acera, Guillermo Palou Márquez, Etna Abad, Laura Ortet, Anna Benito, Yacine Kharraz, Carlos M. Martínez, Elisabetta Mereu, Fran Supek, Ana Janic

## Abstract

Immune checkpoint blockade has transformed cancer therapy, but current biomarkers such as tumour mutation burden often fail to reliably predict clinical benefit. One proposed reason for this discrepancy is the activity of nonsense-mediated mRNA decay (NMD), a cellular quality-control pathway that degrades mutant transcripts, potentially reducing the presentation of neoantigens that would otherwise stimulate anti-tumour immunity. To address this, we identified publicly available small-molecule inhibitors targeting the NMD factors SMG5 and SMG7 (NMDi), which we have prioritise as the NMD components most strongly associated with NMD efficiency across TCGA tumours, and evaluated their therapeutic potential *in vivo*. In a syngeneic, DNA repair deficient mouse model of lung adenocarcinoma, NMDi treatment alone substantially reduced tumour burden, and its combination with anti–PD-1 therapy led to additive benefit either treatment alone. These effects were immune-dependent and specifically required CD8□ T cells. Transcriptomic and single-cell analyses revealed that NMDi reprograms the tumour immune microenvironment, enriching for clonally expanded, cytotoxic CD8□ T cells and altering macrophage states toward those associated with tumour regression. Whole-genome sequencing of tumours revealed that NMDi also promotes immunoediting, leading to negative selection against immunogenic mutations and coding indels, with selective pressure comparable to or greater than that induced by anti–PD-1 treatment. Guided by these results, we generated a new protein language model trained on tumour mutations that can identify neopeptides with immunogenic properties, revealing that the NMDi tumour landscape provides a rich genomic and immunopeptidomic setting for exposing neoantigens that evade immunoediting. Together, these pre-clinical and integrative genomic-transcriptomic insights position NMD inhibition as a promising immunostimulatory strategy that can potentiate immune checkpoint blockade efficacy in tumours.

## Introduction

Immunotherapy, particularly immune checkpoint blockade (ICB), has shown significant efficacy across multiple cancer types^1^. Current patient selection methods rely on several biomarkers such as programmed death-ligand 1 (PD-L1) tumour and immune cell expression^2^, circulating tumour DNA (ctDNA)^3^, microsatellite instability (MSI) status^4^ and tumour mutation burden (TMB) based on the observation that ICB tends to be more effective in hypermutated tumours^5^. In addition, only ∼10% of human tumours display high TMB, and meta-analysis of five cancer types treated with ICB shows that TMB correctly classifies only about 66% of patients^6^. More refined models are challenging to establish due to the lack of a robust correlation between known genomic markers and clinical response^7^.

A substantial fraction of patients predicted to respond based on high TMB exhibit poor outcome, with a false discovery rate exceeding one-third, and by some sources up to a half ^8^. A key factor contributing to this discrepancy was postulated to be the nonsense-mediated mRNA decay (NMD) pathway^9^. NMD silences the expression of mutant mRNAs, preventing translation and presentation of peptides from these proteins on MHC class I complexes and thus potentially suppressing the presentation of neoantigens^6,10^. It appears that only those frameshift (FS) indel mutations that escape NMD predict immunotherapy response, but other indels are not predictive ^10^. Importantly, previous work has shown that inhibition of NMD pathway could have anti-tumour effect and elicit an immune response^11–13^. However, the molecular mechanisms driving the anti-tumour immune response remain to be elucidated. Given the convergence of mechanisms, pharmacological NMD inhibition could potentiate ICB therapy response. Further, the NMD pathway can be inhibited at different points, and implications to efficacy and safety of different interventions are unclear.

Here we evaluated publicly available NMD inhibitors using NMD reporter assays and identified compounds targeting SMG5 and SMG7 (hereafter NMDi) synergistically inhibit NMD activity. These inhibitors exhibited no detectable toxicity *in vitro* or *in vivo*. The combination of NMDi and anti-PD-1 treatment *in vivo*, using an orthotopic murine mismatch repair deficient (dMM) lung adenocarcinoma model, significantly reduced tumour burden, as evidenced by decreased lung weights, compared to either treatment alone, indicating enhanced anti-tumour efficacy. Notably, NMDi alone elicited a strong anti-tumour response in our model, highlighting NMD inhibition as a promising therapeutic strategy for tumours with high TMB. Mechanistically, the therapeutic effect of NMDi depends on immune activation, driven by CD8□ T cells, and involves remodelling the tumour microenvironment to promote clonal expansion of cytotoxic CD8□ T cells. Furthermore, our analysis of tumour evolution WGS data suggest that NMDi therapy results in immunoediting of mutations in the tumour genomes. This indicates a negative selective pressure that reduces neoantigen burden by eliminating highly immunogenic clones, due to NMDi treatment. Therefore, genomic evidence supports that pharmacologically induced NMDi is immunostimulatory, underscoring the co-inhibition of SMG5 and SMG7 as a new approach to treat DNA repair deficient tumours.

## Results

### Pharmacological inhibition of SMG5 and SMG7 synergistically suppresses NMD without inducing detectable toxicity in mice

To prioritise NMD factors for pharmacologic targeting in cancer, we first asked which components of the NMD machinery, when altered by mutation in human tumours, most strongly determine NMD efficiency (NMDeff). We leveraged existing estimates of NMDeff from our recent study^19^ on The Cancer Genome Atlas (TCGA)^15^, which drew on two orthogonal approaches: RNA expression of known, endogenous NMD targeted transcripts, and allele-specific RNA expression of germline nonsense mutations. We associated tumour NMDeff scores with somatic alterations (truncating/missense SNVs and CNA gains or deletions) in 15 NMD genes, including core NMD factors: UPF1, UPF2, UPF3A, UPF3B, SMG1, SMG5–9, and exon-junction complex factors: EIF4A3, CASC3, MAGOH, MAGOHB, RBM8A). Of these factors, the SMG1 kinase that phosphorylates the central NMD factor UPF1 is a common target for various NMDi compounds, ranging from lowly-specific inhibitors of SMG1 such as caffeine and wortmannin, to more specific SMG1i-11 (11j)^17^ and its derivative KVS0001^13^. Further, compounds that selectively inhibit SMG5 (NMDI-1 compound disrupts the SMG5 interaction with phospho-UPF1), SMG7 (NMDI-14 compound disrupts SMG7 interaction with UPF1^20^), and finally EIF4A3^21,22^. We acknowledge that NMD can be inhibited in experiment by “blunt” inhibition of translation (cycloheximide, thapsigargin, tunicamycin) however toxicity would preclude use outside of specific applications (e.g. homoharringtonine^24^)

To prioritise target genes whose alteration would inhibit NMD, we analysed genetic changes across TCGA genomes and transcriptomes. For that we ranked genes within pan-cancer and each TCGA cancer type by association p-value (1 = strongest, 15 = weakest), and an average rank was computed across cancer types. SMG5 showed the top-ranking associations with NMDeff across all 15 NMD-related genes (SNV mutations: mean rank 5.55; CNA gains: 6.26), with SMG7 third-ranking target (mutations mean rank 6.10; CNA gains: 6.39) (Figure 1A). The EIF4A3 is, interestingly, also highly prioritised as a target based on NMDeff association across TCGA tumours, sitting between SMG5 and SMG7 (Figure 1A). In contrast, the commonly used SMG1 inhibitor in various studies was sixth-ranked in our prioritisation. Considering ranking of individual cancer types, the p-value association of gene with NMDeff was significantly stronger for SMG7 and SMG5, compared to (Figure 1B and Supplementary Figure 1A). Further evidence, based on (1) NMD factor mRNA expression levels correlation with same NMDeff in TCGA^19^ (Supplementary Figure 1B), (2) gene essentiality patterns in large databases CRISPR and RNAi screens in DepMap^26,28^ (Supplementary Figure 1C-D), and (3) CIBERSORT estimates of CD8+ T-cell infiltration in TCGA tumours^30^ (Supplementary Figure 1E) similarly prioritised SMG5 and/or SMG7 as strong inhibition targets.

**Figure 1.**
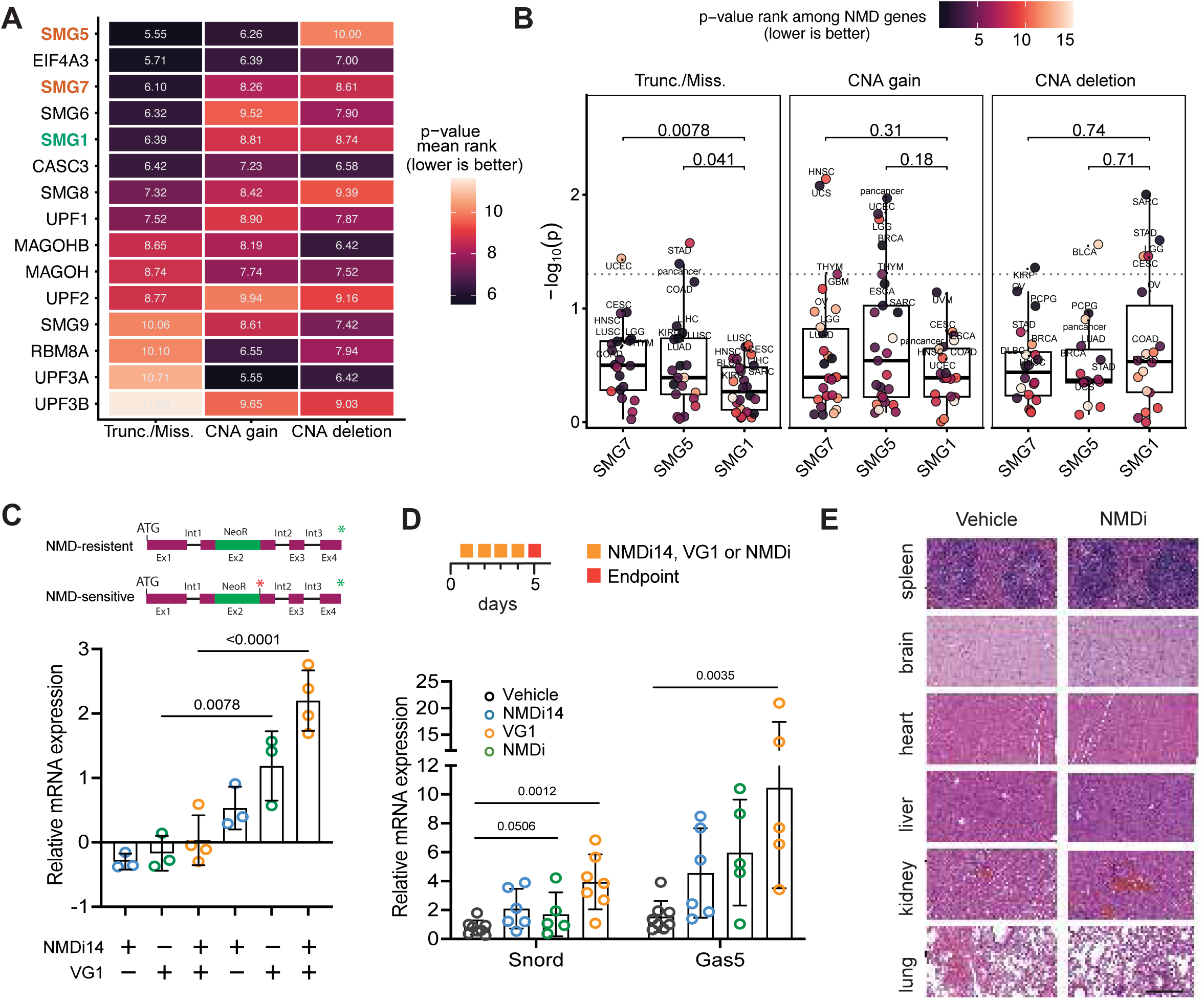
Pharmacological inhibition of SMG5 and SMG7 synergistically suppresses NMD. A) Ranking of 15 core NMD factors (incl. EJC genes) based on the strength of general linear model association between their somatic alterations and tumor-level NMD efficiency (NMDeff) across 31 TCGA cancer types and pan-cancer. The plot displays the mean rank of p-value (lower rank indicates stronger association) derived from linear models associating NMDeff with somatic truncating/missense SNVs, with copy-number alteration (CNA) gains, and with CNA deletions. B) Statistical significance (p-values) of linear model associations between somatic alterations and NMDeff scores for prioritized targets SMG5, SMG7, and SMG1 across cancer types and pan-cancer (other genes visualised in Supplementary Figure 1). Points are coloured according to the gene’s relative rank among 15 NMD factors within that specific cohort (colour gradient: 1 = top rank/strongest association to 15 = lowest rank/weakest association). Statistical significance of the difference in distributions of per-cancer-type p-values between SMG5/7 genes and SMG1 gene as a reference was determined using a one-tailed Mann–Whitney U test. C) Relative quantification of NMD reporter mRNA expression in HeLa cells stably expressing either the NMD-resistant or -sensitive mTCRβ constructs, as depicted in the schematic (top, the red asterisk represents PTC the green asterisk represents the canonical STOP codon). Cells were treated for 6h with 1µM NMD inhibitors (NMDi14, VG1, NMDi) or left untreated. RNA was analyzed by qRT–PCR, and reporter mRNA levels were normalised to relative NeoR mRNA levels. Data represents n=3-4 independent experiments. Data represent mean ± SD. Statistical analysis was performed using one-way ANOVA. D) Relative quantification of NMD endogenous reporter genes mRNA in the ileum of mice treated with 7,5 mg/kg NMDi14, 20 mg/kg VG1, or combination NMDi (hereafter 7,5 mg/kg NMDi14 + 20 mg/kg VG1) for 5 consecutive days, as depicted in the schematic (top). Every dot represents a mouse. Data represents mean ± SD. P values were calculated using one-way ANOVA. E) Representative H&E-stained sections of various organs, as indicated, from mice treated with 5 consecutive doses of NMDi or vehicle. Vehicle (n=1) and NMDi (n=5). Scale bar represents 50µm.

Guided by this prioritisation of SMG5 and SMG7 as leading candidate targets, we next sought to test whether pharmacologic inhibition of these factors functionally suppresses NMD. Thus, we assessed the efficacy of publicly available SMG5 and SMG7 NMD inhibitors, VG1 (NMDI-1 analog)^32^ and NMDI14, using a HeLa cell-based NMD reporter system expressing GFP-tagged TCRβ minigenes that distinguish NMD-sensitive from NMD-resistant transcripts (Figure 1A)^16^. Inhibition of NMD results in upregulation of the NMD-sensitive mRNA containing a premature termination codon (PTC)^14^. We treated HeLa cells expressing these two reporters constructs with either VG1 or NMDI14, for 6 hours. Each compound alone induced the expected increase of the PTC-mutated reporter at low concentrations (1µM). However, combining VG1 and NMDI14 (hereafter NMDi) at a total of 1□μM of NMD inhibitors resulted in a significantly greater upregulation, more than a 2-fold increase, compared to either treatment alone (Figure 1C).

We next evaluated the ability of NMDI14 and VG1, alone or combination to attenuate NMD in mice *in vivo*. We treated wild-type mice with NMDI14 and VG1, or their NMDi combination once daily for 5 consecutive days and quantified the steady-state levels of the NMD endogenous targeted Snord and Gas5 transcripts^18^ in the ileum (Figure 1D). We found that the abundance of both NMD targets was significantly increased relative to untreated controls when treated with NMDi combination therapy.

There have been reports on the importance of NMD in several vital tissues, raising the question of whether an NMD inhibitor would find its therapeutic use in the clinic ^19^. For that we treated wild-type mice with either vehicle or NMDi combination and monitored the acute impact 5 days post-treatment. Histological analysis revealed no gross tissue abnormalities in the major organs in response to NMDi (Figure 1E), consistent with previous findings for certain NMD inhibitors^18,23^. These results support the potential therapeutic application of the SMG5 and SMG7 co-inhibition in the clinic.

### Targeting NMD boosts immunotherapy response in mismatch repair deficient lung tumours *in vivo*

Frameshift mutations in cancer cells with deficient DNA mismatch repair (dMMR) generate PTC-containing transcripts that are negatively regulated by NMD^25,27^. Inhibiting NMD could therefore enhance the accumulation of these tumour specific antigens in tumours rich with frameshifting indels, such as dMMR tumours. Thus, to investigate whether NMD inhibition can improve the response to immunotherapy, we used a syngeneic murine model of dMMR lung adenocarcinoma, recently shown to be robustly sensitive to PD-1 blockade therapy^29^, and therefore a suitable tool for study of ICB potentiating treatments. This model is based on the *Kras^G12D^; p53^-/-^*lung adenocarcinoma cell line^31^ engineered to knock out Mlh1, a key MMR tumour suppressor gene (hereafter Mlh1^KO^)^33,34^.

To test the *in vivo* NMDi therapy and synergistic effect between anti-PD-1 and NMDi, we intravenously injected GFP-NLS tagged Mlh1^KO^ cells to generate orthotopic lung tumours. Seven days post-injection, mice were randomised into four treatment groups: (1) IgG2a isotype or vehicle control, (2) anti–PD-1 therapy, administered as three systemic injections, (3) NMDi treatment consisting of five consecutive daily systemic injections of VG1 and NMDI14, and (4) combination therapy with both anti–PD-1 and NMDi. Treatments were administered over the course of two weeks (Figure 2A). In 5 independent experiments, we demonstrated that the combination of NMDi and anti–PD-1 therapy resulted in superior anti-tumour efficacy, as evidenced by significantly lower lung weights compared to NMDi, anti–PD-1, or control treatments (Figure 2B, Supplementary Figure 2A). Importantly, we observed a significant reduction in the percentage of cancer cells at the experimental endpoint in both NMDi and anti–PD-1 treated mice (Figure 2C, Supplementary Figure 2B). This difference in cancer burden was supported by a substantial effect size (R² = 0.4033, one-way ANOVA), indicating that approximately 40% of the variance in cancer cell percentages was explained by differences among treatment groups. Histopathological analysis further confirmed that tumour burden was markedly reduced in mice treated with either NMDi alone or NMDi and anti–PD-1 (Figure 1D, E). Similarly, the anti-tumour effects of combined NMDi and anti–PD-1 therapy were replicated in similar orthotopic lung cancer model based on the *Kras^G12D^; p53^-/-^* cell line engineered to knock out Pms2, involved in MMR pathway (Supplementary Figure 2C). Overall, these pre-clinical assays demonstrate that combined anti–PD-1 and NMDi treatment effectively enhances tumour cell immunogenicity. Notably, NMDi alone elicited a strong anti-tumour response in our model, highlighting NMD inhibition as a promising therapeutic strategy for DNA repair deficient tumours.

**Figure 2.**
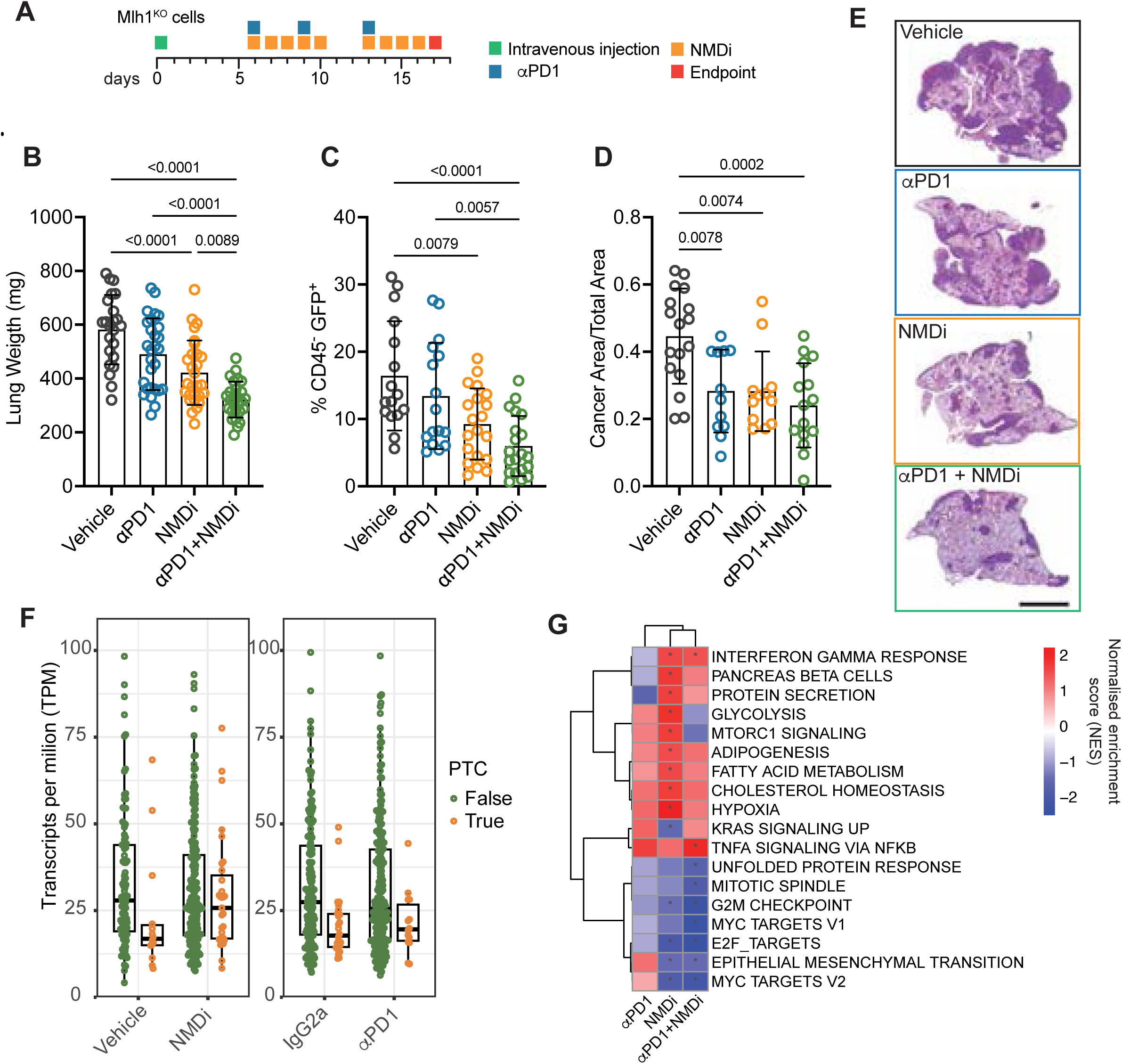
Pharmacological inhibition of SMG5 and SMG7 enhances tumour immunogenicity and potentiates anti–PD-1 immunotherapy. D) Schematic representation of the NMDi and aPD1 administration regime in C57BL/6J mice. E) Wet weight (mg) of the lung of vehicle/IgGs (*n*=23), αPD1 (n=28), NMDi (n=27), or NMDi+ αPD1 (n=28) treated mice across 5 independent experiments. F) Percentage of GFP+CD45- cancer cells of vehicle/IgG2a (n=16), αPD1 (n=16), NMDi (n=20), and NMDi+ αPD1 (n=21) treated mice across 3 independent experiments analysed by flow cytometry. G) Quantification of tumour area relative to lung area from mice treated with vehicle/IgG2a (n=18), αPD1 (n=12), NMDi (n=12), or NMDi+ αPD1 (n=15) across 3 independent experiments. Data represent mean ± SD. One-way ANOVA. Data represent mean ± SD. Every dot represents a mouse at 18 days following i.v. Mlh1^KO^-NLS-GFP cells injection. P-values are calculated using one-way ANOVA. H) Representative images of H&E-stained left lung lobes. Scale bar = 3 mm. I) NMD inhibition stabilises premature-termination-codon (PTC) transcripts. (left) Raw transcripts-per-milion (TPM) values for transcripts carrying at least one annotated PTC (stop-gain or frameshift indel). Only genes whose median raw TPM exceeded 10 in either treatment were analysed. P values were calculated by Mann-Whithey U-test. Vehicle vs NMDi, U=144, p=0.038; IgG2a vs αPD1, U=159, p=0.26. J) Heatmap showing MSigDB Hallmark gene set enrichment analysis (GSEA) in cohorts of mice in indicated conditions, vehicle/IgG2a (n=5), aPD1 (n=4), NMDi (n=4), or NMDi+ αPD1 (n=3). Rows include every pathway that reached false-discovery significance (p adjusted value < 0.05) in at least one comparison. Asterisks denote pathway contrast pairs with p adjusted value < 0.05.

To confirm that NMDi impairs degradation of PTC-sensitive transcripts (those that bear nonsense or frameshift mutations) *in vivo*, RNA-Seq of sorted GFP+ tumour cells showed significant upregulation of these transcripts in NMDi-treated tumours compared to controls (Figure 2F), indicating effective NMD inhibition. As expected, anti-PD-1 treatment had no such effect, supporting that transcript stabilisation of PTC-bearing transcripts is specific to NMDi.

To investigate the molecular effects of PD1 blockade, NMDi, and their combination, we performed RNA-Seq of sorted GFP+ tumour cells and revealed that anti-PD-1 and NMDi monotherapies and the combination therapy induce broadly related gene signatures, including shared upregulation of pro-inflammatory TNFa signalling and interferon-γ signalling specifically enriched in NMDi-treated tumours. The epithelial-to-mesenchymal transition gene program was suppressed with NMDi treatment, in contrast to anti-PD-1 only treatment. NMDi also downregulated proliferative pathways (e.g., MYC targets, E2F targets, G2M checkpoint), supporting its cytostatic effects, and this effect was further enhanced in the combination treatment. These results highlight NMDi’s antiproliferative and immunostimulatory actions and its synergy with PD1 blockade (Figure 2G).

Together, these preclinical studies show that combining anti–PD-1 therapy with NMD inhibition significantly enhances tumour immunogenicity by promoting both immune stimulating and cytostatic effects. Notably, NMDi therapy alone also elicited a strong anti-tumour response in our model, highlighting its potential as an independent cancer therapeutic strategy driven by immune activation mechanisms.

### CD8□ T cells drive NMD-mediated anti-tumour activity

To assess whether the improved outcomes observed in mice treated with the NMD inhibitor were dependent on the immune response, we injected dual luciferase/GFP-labelled Mlh1^KO^ lung cancer cells into both wild-type C57BL/6 immunocompetent (BL/6) and immunodeficient NOD-SCID Gamma (NSG) host mice, followed by systemic NMDi treatment (Figure 3A). Bioluminescence imaging (BLI) revealed a strong anti-tumour response in immunocompetent mice, whereas no significant effect was observed in immunodeficient hosts (Figure 3B-D). These results indicate that the therapeutic efficacy of NMDi requires a functional immune system.

**Figure 3.**
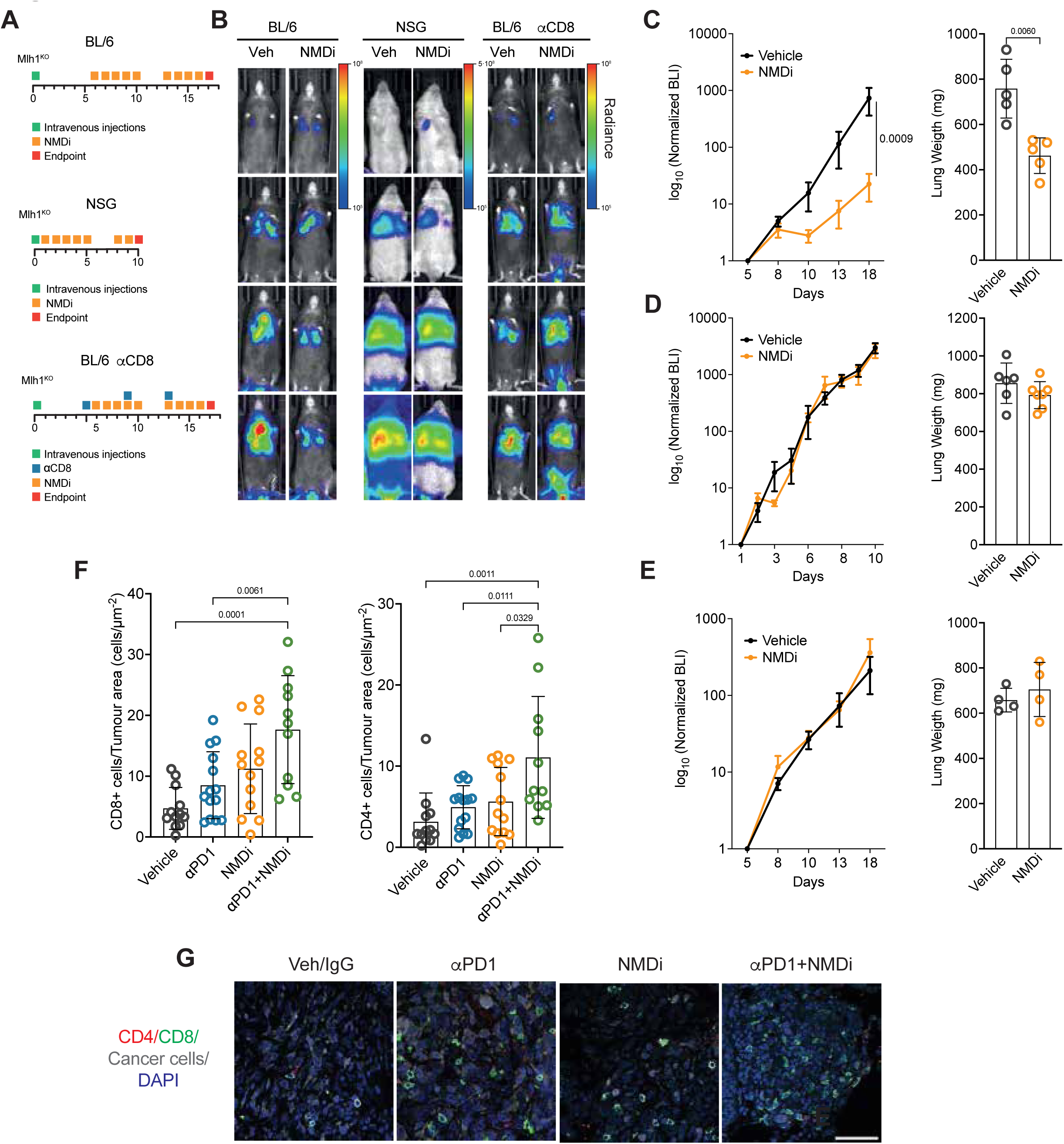
NMDi-mediated anti-tumour immunity relies on CD8^+^ T cells. A) Schematic representation of the NMDi and αCD8 administration regime in C57BL/6J (BL/6) and NSG mice. B) Representative bioluminescence (BLI) images of indicated cohorts. Colour scale indicates intensity of radiance. C-E) Growth curves (Left) and wet weight of the lungs (right) of mice treated with the vehicle (N=4-6) or NMDi (N=5-7). Data for growth curves represent mean ± SEM in log_10_ scale, p values are calculated by two-way ANOVA. Data lung weights (mg) represent mean ± SD, p-values are calculated by two-tailed Student’s t-test. F) Quantification of CD8^+^ T (right) or CD4^+^ T (left) cells per tumour area in lung from mice treated with vehicle/IgG2a (n=4), aPD1 (n=4), NMDi (n=4), or NMDi+ αPD1 (n=4) At least 2 tumour lesions were analysed per condition in 3 independent experiments, each dot represents a tumour lesion. Data represent mean ± SD. P values are calculated by one-way ANOVA. G) Representative images of CD4 and CD8 immunostaining in indicated conditions. Cancer cells are labelled in grey, CD8^+^ T cells green, CD4^+^ T cells in red, DNA in blue. Scale bar represents 5mm.

T cell infiltration into tumours is a critical determinant of response to immunotherapy, as it reflects the immune system’s ability to recognise and attack cancer cells^35^. To analyse the role of immune cells, specifically T cells, in NMDi therapy, we performed antibody-mediated CD8□ T cell depletion studies in mice bearing orthotopic Mlh1^KO^ lung tumours treated with systemic NMDi (Figure 3A). Successful CD8□ T cell depletion was confirmed by flow cytometric analysis of lung tissue (Supplementary Figure 3A, B). This depletion completely abrogated the antitumour efficacy of NMDi therapy (Figure 3E), demonstrating that CD8□ T cells are essential mediators of NMDi-induced anti-tumour immunity. To confirm T cell infiltration at tumour sites, lungs from tumour-bearing mice were collected following NMDi treatment and analysed by immunofluorescence. The results confirmed high levels of CD8^+^ and CD4^+^ T cell infiltration within the tumour sites (Figure 3 F, G), demonstrating active immune engagement at the tumour site.

These findings demonstrate that the anti-tumour efficacy of NMD inhibition is immune dependent and critically relies on functional CD8□ T cells. The absence of therapeutic benefit in immunodeficient hosts, coupled with the loss of efficacy following CD8□ T cell depletion, highlights the essential role of adaptive immunity in mediating NMDi driven tumour control.

### NMD inhibition remodels the tumour immune microenvironment and expands clonally active cytotoxic CD8□ T cells

To characterise tumour infiltrating immune cell populations in dMMR lung tumours following NMDi, we performed single-cell RNA sequencing (scRNA-seq) coupled with single-cell TCR sequencing (scTCR-seq). We profiled 71,977 CD45□ immune cells FACS-isolated from tumour-bearing lungs of immunocompetent mice intravenously injected with GFP-NLS-tagged Mlh1^KO^ cells. Mice were treated with either NMDi or vehicle over the course of two weeks (Figure 2A). Single-cell RNA-seq resolved all major immune compartments and revealed NMDi dependent shifts in their abundance (Figure 4A, B). NMDi treatment generally reduced total T cell frequency yet produced a small, highly cytotoxic CD8□ subset, depleted early/memory B cell and plasma cell pools, and increased hi-ribosome B cells and monocytes (Figure 4B). Within the three tumour-associated macrophage (TAM) states, TAM2 predominated in untreated lesions, TAM1 was enriched in responding tumours, and TAM3 accumulated exclusively in poor responders (Figure 4B). Stratifying NMDi treated samples by outcome (Supplementary Figure 2A) exposed a gradient of lymphoid re-composition, the bad NMDi responder showed broad T cell loss. Good NMDi responders gained precursor and terminally exhausted CD8□ subsets, along with proliferating T cells and NK cells. The very good NMDi responder was distinguished by expansion of precursor exhausted CD8□ T cells and cytotoxic CD8□ T cells, correlating with maximal tumour regression (Figure 4B).

**Figure 4.**
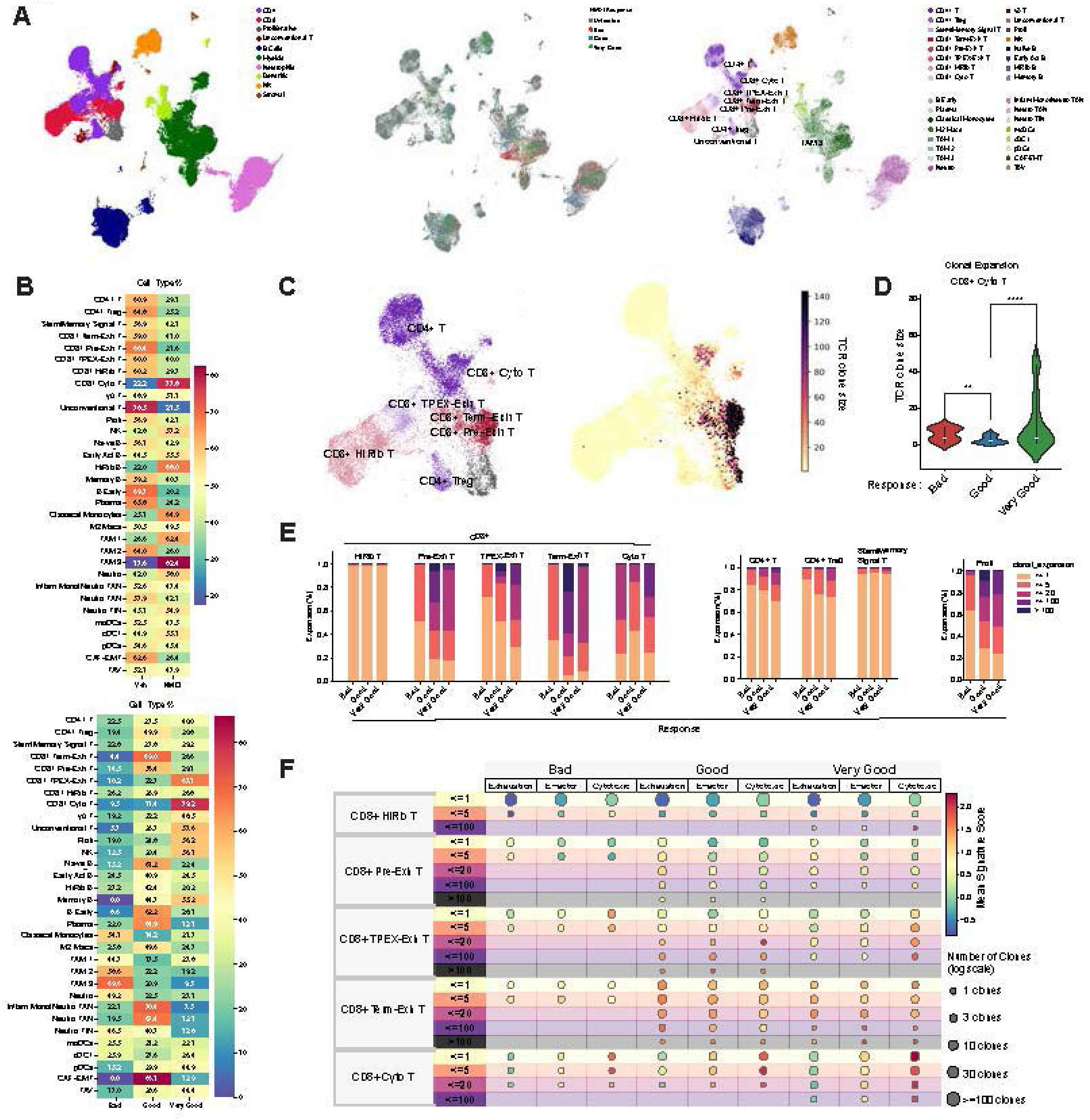
Single-cell profiling uncovers immune remodeling and clonal cytotoxic T cell expansion following NMD inhibition. A) Uniform Manifold Approximation and Projection (UMAP) of 71,977 cells from scRNA-seq samples. (Left) UMAP coloured by broad cell type annotation. Cell types include: CD4 T cells (CD4), CD8 T cells (CD8), B cells (B Cells), Myeloid cells (Myeloid), Neutrophils (Neutrophils), Dendritic cells (Dendritic), Natural Killer cells (NK), Stromal cells (Stromal), Proliferative cells (Proliferative), and Unconventional T cells (Unconventional T). (centre) UMAP coloured by sample condition. “Untreated” refers to vehicle control samples; “Bad”, “Good”, and “Very Good” indicate increasing levels of NMDi treatment response. (right) UMAP coloured by fine-grained cell type annotation. Labels include: CD4+ T cells (CD4+T), regulatory T cells (CD4+Treg), stem/memory-like signaling T cells (Stem/Memory Signal T), terminally exhausted CD8 T cells (CD8+ Term-Exh T), precursor exhausted CD8 T cells (CD8+ Pre-Exh T), TPEX CD8 T cells (CD8+ TPEX-Exh T), high-ribosome CD8 T cells (CD8+ HiRib T), cytotoxic CD8 T cells (CD8+ Cyto T), gamma delta T cells (γδ T), unconventional T cells ( Unconventional T), proliferative cells (Proli), NK cells (NK), naïve B cells (Naïve B), early activated B cells ((Early Act) B), high-ribosome B cells ((HiRib) B), memory B cells ( Memory B), early B cells (B Early), plasma cells (Plasma), classical monocytes (Classical Monocytes), M2-like macrophages (M2 Macs), tumor-associated macrophages 1–3 (TAM 1–3), neutrophils (Neutro), inflammatory monocytes/neutrophils (Inflam Mono/Neutro TAN), tumor-associated neutrophils (Neutro TAN), tumor-infiltrating neutrophils (Neutro TIN), monocyte-derived dendritic cells (DC(moDCs)), conventional dendritic cells type 1 (cDC1), plasmacytoid dendritic cells (pDCs), cancer-associated fibroblasts with EMT features (CAF-EMT), and tumor-associated vasculature (TAV). B) Heatmaps of cell type proportions across conditions. (Top) Heatmap showing the average proportion of each cell type from untreated (vehicle) and treated (NMDi) samples. (Bottom) Heatmap showing the average cell type distribution across responder categories within the treated samples. C) UMAP of T cell subset (19,357 cells). (Left) UMAP colored by fine-grained T cell subtype annotations. (Right) UMAP colored by clonotype size. D) Violin plot showing clonotype size across treatment responder categories for cytotoxic CD8 T cells. p-values were calculated using a two-sided Mann–Whitney U test. Asterisks denote statistical significance: P < 0.01 (**), P < 0.0001 (****). E) Clonotype expansion and response analysis. Barplot showing the distribution of cells per T cell subtype across clonotype expansion categories. E) Dot plot summarizing signature scores across CD8+ T cell subsets and clonotype categories. Signature scores for exhaustion, effector, and cytotoxic profiles are shown across CD8+ T cell subtypes and clonotype expansion categories, stratified by treatment response.

Joint single-cell TCR analysis pinpointed how clonotypic dynamics paralleled these transcriptional changes (Figure 4C–F). Terminally exhausted CD8□ T cells carried the largest clones in all conditions, suggesting disease driven expansion that persisted despite NMD inhibition (Figure 4C, Supplementary Figure 4A). In contrast, the very good NMDi responder displayed moderate clone sizes overall but a markedly higher proportion of expanded clones in precursor exhausted CD8□, cytotoxic CD8□, CD4□ stem-memory and proliferating T cells, cytotoxic CD8□ expansion was significantly greater than in all other groups (Figure 4D, Supplementary Figure 4B,C). Functional gene-set scoring reinforced this pattern (Figure 4E). Cytotoxic CD8□ cells from the very-good responder were simultaneously highly cytolytic (Prf1, Gzms), minimally exhausted (low Pdcd1, Ctla4) and clonally expanded, whereas precursor exhausted CD8□ T cells in good NMDi responders retained cytotoxic potential but began to acquire exhaustion markers, and bad responders showed uniformly low expansion and close to muted effector or cytotoxic signatures (Figure 4D).

Together, these data indicate that successful NMDi therapy remodels the immune milieu towards a focused pool of clonally expanded, cytotoxic-competent CD8□ T cells while selecting TAM states associated with favourable tumour control.

### NMD inhibition induces immunoediting in cancer genomes

To assess whether NMD inhibition alters immune-driven selection, we performed whole genome sequencing of FACS purified GFP+ tumour cells, followed by single nucleotide variant (SNV) and indel calling and analysis. Immune dN/dS ratios^36^ were then calculated by comparing nonsynonymous to synonymous mutations in predicted MHC class I presented regions versus non-antigenic regions across tumours from control, anti-PD-1, NMDi, and combined NMDi and anti–PD-1 treatment groups. Control tumours showed immune dN/dS values clustering around zero, consistent with neutral evolution. In contrast, all three treatment groups showed a consistent downward shift in immune dN/dS values (median < 0), indicating stronger purifying selection against immunogenic mutations. This effect was most pronounced in NMDi and combined NMDi and anti–PD-1 treated tumours, where many samples fell below the neutral threshold (Figure 5A). Across different epitope prediction thresholds (NetMHCpan percentile rank < 0.5% or 1%), rare examples of tumours occasionally showed immune dN/dS values above 1, likely due to sampling noise or low mutation counts. NMDi-treated tumours consistently exhibited reduced immune dN/dS values (medians = 0.47–0.97 across different methodology settings; rate-ratio vs. control = 0.66–0.99), suggesting up to ∼53% of potentially immunogenic missense mutations are eliminated under NMDi treatment, while by the same estimate up to ∼29% would be removed by negative selection without treatment. Remarkably, the effect of NMDi on immunoediting was similar or stronger than that of anti-PD-1 monotherapy (median = 0.73–1.01; rate-ratio = 0.67–1.05). The combination therapy did not further increase the negative selection on MHC-presented peptides (median = 0.55–0.98; rate-ratio = 0.60–1.06) (Figure 5A).

**Figure 5.**
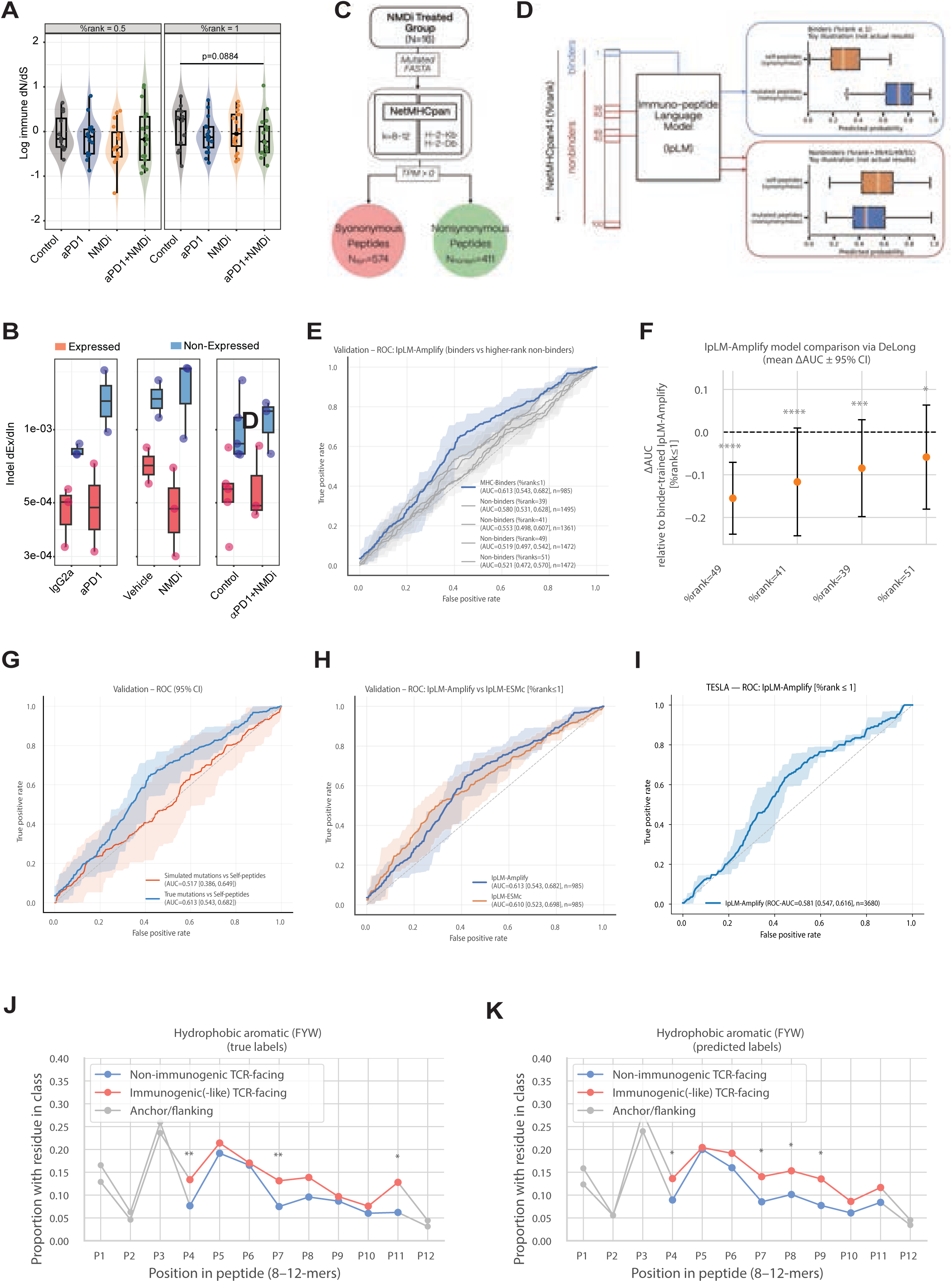
NMDi enhances immune-mediated negative selection against tumour neoantigens. A) Immune dN/dS indicates stronger immune-mediated negative selection in genomes of treated tumours. Log-transformed immune dN/dS values are shown for tumours treated with Control (IgG), αPD1, NMDi or the αPD1+NMDi combination. Each point represents one tumour sample. The horizontal dashed line marks the neutral expectation (log dN/dS = 0). Left panel includes mutations ranked within the top 0.5% binders predicted by NetMHCpan4.1 (%rank = 0.5). Right panel represents the broader set of immune genes with peptides in the top 1% bindiers (%rank = 1). All results here are limited to genes with detectable expression (TPM > 0). B) Exonic to intronic indel counts ratio (dEx/dIn). Indel dEx/dIn values computed as (exonic+1)/(intronic+1), are shown for three pairwise comparisons: IgG2a vs αPD1 (left panel), Vehicle vs NMDi (center), and Control (pooled IgG2a + Vehicle) vs αPD1+NMDi (right). Within each panel, genes are stratified by transcriptional status, expressed genes (TPM > 0) denoted in red, and non-expressed in grey blue. Each point represents one mouse. C) Schematic overview of the selection of immunogenic-like peptides and matched self-peptide controls, both with predicted high-affinity MHC binding. D) Illustration of the sampling rationale for the training dataset (blue) where the IpLM is trained on the %rank ≤ 1 MHC binder set; within this set of MHC binders class separation between nonsynonymous (positive) and synonymous (negative) peptides is expected. As an additional control experiment, within peptides sampled from various non-MHC-binder bins (%rank = 39/41/49/51), no discriminative signal should arise between mutated and synonymous peptides. Boxplots in this panel are cartoons used solely to illustrate the expected model behavior. E) ROC curves comparing IpLM-Amplify trained on positive and negative examples sampled from high-affinity MHC binders (%rank ≤ 1), compared with control models trained on positives and negatives from non-binder peptide strata (%rank = 39/41/49/51). F) DeLong statistical test for AUCs difference between the binder-trained model and the control, non-binder–trained variants. Points denote the mean ΔAUC across cross-validation folds (relative to the binder-trained model), with horizontal bars showing the 95% C.I. G) Performance comparison of IpLM-Amplify model trained on observed neoantigens (nonsynonymous mutations), versus control models trained on simulated neoantigens generated in the same genes according to SBS96 mutational signatures, with transcription strand bias information preserved (i.e. SBS192). H) Direct comparison between IpLM-Amplify and IpLM-ESM-C-, two different foundation language models, both fine-tuned on the positives and negatives from the high-affinity binder set (%rank ≤ 1). I) ROC curve showing the performance of IpLM-Amplify when evaluated on the TESLA neoantigen benchmark dataset. The curve represents the mean ROC across cross-validation folds, with shaded bands indicating the 95% confidence interval. J) Hydrophobic enrichment analysis using observed mutation labels (“true” labels), comparing peptides generated by nonsynonymous versus synonymous mutations. Red and blue curves show residue frequencies at TCR-facing (non-anchor) positions for immunogenic-like and self-like peptides, respectively, while grey curves indicate anchor and flanking positions. Asterisks denote positions with significant associations between peptide class and aromatic residue presence (*p < 0.05, **p < 0.01; Cochran–Mantel–Haenszel test stratified by peptide length). K) Same analysis as in J, but using IpLM-Amplify classifier–predicted labels for the same NMDi-derived high-affinity binder peptides, allowing us to probe how the classifier partitions nonsynonymous (immunogenic-like) versus synonymous (self-like) peptides.

Default analyses were performed while restricting to only expressed genes (median TPM > 0 in each group, Figure 5B), however the relative effects of stronger negative selection upon treatment remain qualitatively even if no expression filtering is performed (Supplementary Figure 5 A, B). These findings indicate immunotherapy-induced immunoediting and indicate that NMDi triggers comparable or greater selective pressure on immunogenic mutations than the PD1 blockade does in this lung cancer model.

To extend this observation beyond point mutations, we analysed indels, previously linked to immunotherapy response and NMD sensitivity^37,38^, by calculating the ratio of coding (dEx) to intronic (dIn) indels per tumour (Figure 5B). Lower dEx/dIn ratios indicate stronger negative selection against potentially immunogenic coding indels. Anti-PD-1 treated tumours showed a modest drop in this ratio relative to their matched IgG2a controls, while NMDi-treated tumours displayed a more pronounced reduction versus the matched vehicle treated controls. The strongest signal was seen in expressed genes (median TPM > 1), mirroring the SNV-based results.

Alternative splicing can generate immunogenic neopeptides^39,40^, and we investigated whether NMDi could restore expression of such events to enhance tumour clearance. NMDi treated tumours showed significantly more intron retention events than matched controls, indicating that these transcripts are typically degraded by NMD, unlike in anti–PD-1–treated tumours (Supplementary Figure 6A). No substantial changes were observed in exon skipping or alternative 3’ or 5’ splice site usage (Supplementary Figure 6B–D). However, the ΔPSI distribution for retained introns was significantly shifted (p = 0.0051; padj = 0.0135), and a modest but significant shift was also seen for skipped exons (p = 0.0067; padj = 0.0135), while changes in A3 and A5 usage were not significant (p > 0.197; padj > 0.26). These results suggest that NMDi selectively promotes intron retention, and to a lesser extent exon skipping, distinguishing it from anti–PD-1 treatment.

Finally, we assessed positive selection in individual genes using the dNdScv tool. Although many candidate genes were shared across groups, likely due to mutations acquired during *in vitro* expansion, differences in gene-level significance profiles emerged between treated and control tumours (Supplementary Figure 5C). Treated tumours showed improved q-values for several genes, suggesting treatment-associated enrichment of mutations under positive selection. Notably, the similarity between NMDi and anti-PD-1 treated group driver gene profiles further supports overlapping selective pressures, consistent with a shared mechanism of immune stimulation by NMDi and anti-PD-1.

These various genomic evolutionary analyses demonstrate that pharmacological NMD inhibition increases immune-driven negative selection against both point mutations and indels in dMMR tumours, comparable to or greater than genomic effects PD-1 blockade, reinforcing its potential as a standalone therapeutic or an adjuvant to immunotherapy.

### Protein language model trained on NMDi treated cancer genomes capture immunogenicity signals beyond MHC binding

The immunoediting signatures described above relied on predictions of MHC binding, yet antigen presentation represents only one step in immune recognition. Many MHC-presented peptides fail to elicit T-cell responses because TCR recognition imposes additional, poorly characterized constraints on immunogenic sequences^45,46^. We hypothesised that protein language models (neural networks pre-trained on large protein datasets) could be fine-tuned on mutations from NMDi-treated tumors to capture these complementary immunogenicity features. For that we developed Immuno-peptide Language Models (IpLMs) by fine-tuning the Amplify foundation model^47^ and ESM Cambrian (ESM-c)^48^ on peptide sets derived from our somatic mutation data from NMDi treated tumours (Figure 5C). Critically, both training classes, nonsynonymous mutation-derived neoantigens (positives) and synonymous mutation-derived self-peptides (negatives), were restricted to high-affinity MHC binders, ensuring that any discriminative signal could not reflect MHC binding itself (Figure 5C, D). IpLM-Amplify achieved a ROC-AUC of 0.613 on held-out data, comparable to IpLM-ESM C with ROC-AUC of 0.610 (Figure 5 H). Control experiments for IpLM-Amplify confirmed this reflected genuine immunogenicity, models trained on non-MHC-binders showed significantly lower performance (p = 0.012 to 1.6×10□□) (Figure 5E, F), while training on simulated mutations preserving mutational signatures but lacking biological selection nullified performance entirely (AUC = 0.517) (Figure 5G). Fine-tuning of IpLM-Amplify induced a modest but consistent reorganization of the embedding space for both classes, nudging the internal representations of the neural net to opposite directions (Supplementary Figure 7E, F).

To further understand the biological basis of these predictions, we examined position-specific amino acid composition. As expected from MHC binding constraints, anchor positions (P2 and C-terminus) showed identical residue usage between immunogenic and non-immunogenic peptides. However, within the TCR-facing region, immunogenic peptides displayed significant enrichment of aromatic residues (F/Y/W) at position P4 and P7 (Figure 5J), with aliphatic hydrophobic residues (L/I/V/M) similarly enriched at P4, P6 and P8 (Supplementary Figure 7 A). These patterns align with structural evidence that aromatic and hydrophobic residues at TCR-contact positions stabilize the trimolecular peptide-MHC-TCR complex^49–51^. Reassuringly, IpLM-Amplify recapitulated the aromatic enrichments, especially at P4 and P7 (Figure 5K), indicating the model learned functionally relevant sequence features rather than artefacts. In contrast, the corresponding aliphatic pattern was not recovered (Supplementary Figure 7C). As a negative control, charged residues (D/E/H/K/R) showed no significant enrichment or depletion at TCR-facing positions in either the training data or model predictions (Supplementary Figure7B, D), consistent with prior structural studies^50^.To assess performance beyond murine NMDi tumour dataset, we evaluated IpLM-Amplify on the independent TESLA neoantigen benchmark from human tumours. The model achieved an above-random AUC of 0.581 (95% CI: 0.547–0.616), demonstrating that immunogenicity-relevant sequence features can transfer across the species and other biological differences (Figure 5 I).

Together, these findings demonstrate that the NMDi tumour dataset creates a uniquely informative genomic and immunopeptidomic landscape for learning determinants of neoantigen immunogenicity and provides a powerful training ground for advancing neoantigen prediction for therapeutic targeting.

## Discussion

Our study adds to the growing body of evidence supporting NMD as a tractable therapeutic target in oncology^11,25^. Mechanistically, our approach selectively targets the downstream NMD effectors SMG5 and SMG7, using two small molecules, VG1 and NMDI14. This dual inhibition disrupts SMG5- and SMG7-dependent mRNA degradation resulting in robust and synergistic NMD suppression with minimal toxicity.

Here we demonstrate that NMDi enhances anti-tumour immune responses in dMMR lung cancer. We show that NMD inhibition leads to improved tumour control, particularly when combined with PD-1 immune checkpoint blockade. This combinatorial approach significantly reduces tumour burden *in vivo* compared to either monotherapy, indicating a synergistic therapeutic benefit. Importantly, the anti-tumour effect of NMD inhibition was abrogated by CD8□ T cell depletion, confirming that cytotoxic T cells are essential mediators of this immune-driven response.

To understand how NMD inhibition shapes the tumour immune landscape, we performed single-cell RNA and TCR sequencing of immune cells from orthotopic dMMR lung tumours treated with NMDi. This revealed a striking remodelling of the tumour immune microenvironment. While NMDi treatment reduced the overall frequency of T cells, it promoted the emergence of a small but highly cytotoxic CD8□ T cell subset and altered the composition of TAMs, enriching TAM1 in responders and depleting TAM3, which accumulated in non-responders. Notably, successful NMDi therapy was associated with a focused expansion of precursor exhausted and cytotoxic CD8□ T cells, along with proliferating CD4□ stem-like and NK cells. TCR analysis further demonstrated that cytotoxic CD8□ T cells in the best NMDi responders were clonally expanded, suggesting active tumour engagement and immunological memory. These results position NMD inhibition as a modulator of the intratumoral immune composition, driving clonal expansion of effector cells and immune states consistent with effective anti-tumour immunity.

A key mechanistic insight from our study is that NMD inhibition promotes tumour immunogenicity by enabling the accumulation of aberrant, neoantigen-bearing transcripts, which in turn triggers selective immune pressure. This was evidenced by reduced immune dN/dS ratios and depletion of coding indels in NMDi-treated tumours, suggesting that NMDi promotes immunoediting by enhancing the visibility of tumour clones to the immune system. Fine-tuning a protein language model on peptide sets derived directly from somatic mutations and constrained to high-affinity MHC binders, allowed us to recover a measurable immunogenicity signal. By contrasting nonsynonymous-mutation derived peptides with synonymous controls in the NMD-inhibited context, IpLM captures features associated with immunogenicity that are independent of MHC binding strength. This suggests that NMDi-treated samples contain neoantigens that escape complete immunoediting but retain detectable immunogenic potential. Within the broader context of NMDi as an immunostimulatory strategy, our observations support the idea that NMD inhibition can broaden the repertoire of immunologically visible neoantigens. These findings position NMD inhibition as an active driver of immune pressure and clonal selection, reinforcing its potential as a powerful immunostimulatory strategy.

Taken together, our study provides a strong preclinical rationale for the clinical development of SMG5 and SMG7 targeted NMD inhibition strategies, particularly in the context of immune checkpoint therapy. Given the conserved role of NMD and the prevalence of frameshift mutations in hypermutated cancers, we anticipate that the immunostimulatory effects observed here will extend beyond DNA repair deficient lung cancer to other high mutation burden tumours. Future work should explore combination strategies with additional immunotherapies and investigate predictive biomarkers of NMDi sensitivity to enable patient stratification.

## Methods

### Animal experiments

All animal experiments complied with ethical regulations for animal research and were approved by the Ethics Committee for Animal Experiments (CEEA-PRBB, Barcelona, Spain). Animals were euthanised at either the study endpoint or a humane endpoint. Male C57BL/6J and NOD.Cg-Prkdc^scid^ Il2rg^tm1Wjl^/SzJ (NSG) mice were used. Mlh1^KO^ cancer cells were injected into the tail vain to generate an orthotopic syngeneic model of lung adenocarcinoma. Tumour engraftment in the lung occurred by day 6 in C57BL/6J mice (as previously reported ^29^) and by day 1 for NSG. C57BL/6J mice were injected with 5-7·10^5^ NLS-GFP or Luc-GFP labelled Mlh1^KO^ cells. Tumour growth was monitored by BLI imaging once per week, and data were collected using Live Image v.4.3.1 in a Perkin Elmer Living Image system. For in vivo treatments, NMDi14 and VG1 were resuspended in a solution of 5% DMSO (Sigma-Aldrich, D8418-250ML), 20% Cremophor® EL (Sigma-Aldrich, 238470-1SET), and 75% saline (Ern, 999791.5). NMDi14 was prepared at 1 mg/ml and VG1 at 2 mg/ml, and both were administered intraperitoneally at 7.5 mg/kg and 20 mg/kg, respectively. Antibody treatments included 100 µg IgG2a (Leinco Technologies, R1367), 100 µg anti–PD-1 (Leinco Technologies, P372), and 250 µg anti–CD8 (Abyntek Biopharma InVivoMAb, BE0117).

### Cell lines and culture conditions

Murine lung adenocarcinoma cells Kras^G12D^-p53^null^ (KP92) were derived from Kras^LSL-G12D^; Trp53^flox/flox^ mice (gift from Dr. Kate Sutherland, WEHI, Australia). All KP92-derived cell line were grown in 5% CO□ at 37□°C. KP92 were grown in Dulbecco’s Modified Eagle’s Medium/nutrient mix F-12 (DMEM/F12) (ThermoFisher scientific, 11320033) supplemented with 4ug/ml hydrocortisone (Sigma-Aldrich, H4001), insulin-transferrin-selenium mix 1x (Sigma-Aldrich, 41400045), 10% FBS (Gibco, Invitrogen, F9665) and 1% Penicillin-Streptomycin (labclinics, FBS-16A). HeLa were grown in Dulbecco’s Modified Eagle’s Medium (DMEM)-glutamax (labclinics, L0102-500) supplemented with 10% FBS (Gibco, Invitrogen) and 1% Penicillin-Streptomycin (labclinics, FBS-16A). Cells were routinely checked for *Mycoplasma* and all were negative. Mlh1^KO^ cell line was generated as described in ^29^. To generate cells expressing luciferase-GFP and NLS-GFP, we used the lentiviral vector pLEX-hFL2iG (gift from Dr. Antoni Celià-Terrassa, Hospital Research Mar), and pTRIP-SFFV-EGFP-NLS (gift from Dr Sara Sdelci, Centre for Genomic Regulation).

### NMD reporter assays

To test the efficiency of the NMD machinery, HeLa cells were transfected for 24 h with Lipofectamine 2000 (ThermoFisher Scientific, 11668027), according to the manufacturer’s instructions with either 200 ng of minigenes mTCRβ-WT (NMD-resistant) or mTCRβ-PTC (NMD-sensitive). Cells were treated with the selected compounds for 6h at 37°C in 5% CO_2_. mRNA was extracted following TRIzol protocol (ThermoFisher, 15596026) and retrotranscribed with SuperScript IV (ThermoFisher, 18090010) using a gene-specific primer according to the manufacturer’s instructions. The gene specific primer: 5’-AGGCATGCAAGCTTCAGAAC-3’ retrieves the mTCRβ as gene of interest and NeoR as normalizing gene. To test the inhibitors efficiency in vivo, mice were treated with either vehicle, NMD14, VG1 or both NMD inhibitors over 5 days. Ilium was collected, washed in PBS 1x and homogenized in 1 ml of TRIzol to extract total mRNA.

### Tissue processing

Lungs were harvested, weighted, minced and placed into gentleMACS C tubes (Miltenyi Biotec, 130-093-237) containing 3 ml of PBS 1x, 5 % FBS, and 2 mg/ml collagenase A type IV (Sigma-Aldrich, C4-22-1G) for lung tissues. The lungs were carefully minced using the gentleMACS Tissue Dissociator (Miltenyi Biotec, 130-096-427) and lungs were digested for 40 min at 37 °C under continuous rotation and then filtered through 70 μm cell strainers. The filter was then washed by adding 5 ml of complete DMEM twice, and the digested tumours were incubated with 0.5–1 ml of RBC lysis buffer (Merck, 11814389001) for 2 min at 4 °C, then washed with 5 ml PBS containing 10 % FBS and 0.1 % sodium azide (PSA buffer). Tumour pellets were resuspended in PSA to obtain single-cell tumour suspensions to proceed with flow cytometry staining or sorting.

### Flow cytometry and cell sorting

Single cell suspensions were resuspended in PSA and incubated for 30 min at 4 °C with 0.5 μg/1 × 10^6^ cells of anti-CD16/32 antibody (Biolegend, 101302). After washing with PSA, cells were incubated with surface marker-specific antibodies (0.2–1 μg of antibody per 1 × 10^6^ cells, upon titration) for 30 min in the dark at 4 °C. After staining, cells were washed with PSA. The antibodies used can be found in XXX. Flow cytometry analysis and fluorescence-activated cell sorting (FACS) were performed with a Cytek Aurora Spectral Cytometer 5 Laser: 16UV-16V-14B-10YG-8R. For cell sorting samples were stained with aCD45 surface marker-specific antibodies and filtered through a 30 μm nylon mesh (Thermofisher, 352235) BD Influx cell sorter (BD Biosciences, Switzerland) with a stable temperature of 4°C, equipped with 325 nm, 457 nm, 488 nm, 561 nm and 633 nm laser and CD45^+^ GFP- or CD45^-^ GFP^+^ according to the experiment setup. Data analysis was done using the SpectroFlo version 3.3.0 (Cytek Biosciences).

### gDNA and mRNA purification

Sorted cancer cells were retrieved in PBS1x supplemented with 10% of FBS and processed freshly straight after cell sorting. gDNA and mRNA was extracted from as little as 25 thousand cells. gDNA was purified using QIAamp UCP DNA Micro Kit (Qiagen, 56204). Total mRNA was purified using TRIzol protocol.

### Isolation or depletion of CD45+ leukocytes

For scRNA-seq of the immune cell compartment single cell suspensions of lungs were magnetically labelled with CD45 MicroBeads (Miltenyi Biotec, 130-052-301) and passed through a MS column (Miltenyi Biotec, 130-042-201) to either isolate or deplete CD45-expressing cells.

### qRT-PCR analysis

qRT-PCR was performed amplifying the resulting cDNA of the retrotranscription of 10-25 ng of mRNA. The following primers were used: mTCRβ, forward 5’-GCGGTGCAGAAACGCTGTA-3’, reverse 5’-TGGCTCAAACAAGGAGACCTT-3’; NeoR, forward 5’-GCTCGACGTTGTCACTGAAG-3’, reverse 5’-GCAGGAGCAAGGTGAGATGAC-3’; Snord, forward 5’-GCCAGGCCTGTTCAATTTTA-3’, reverse 5’-TGCCTGAGATTTGTCACCAG -3’; Aft4, forward 5’-CACAACATGACCGAGATGAG-3’, reverse 5’-CGAAGTCAAACTCTTTCAGATCC -3’; Gapdh, forward 5’-TTCCAGTATGACTCCACTCACGG-3’, reverse 5’- TGAAGACACCAGTAGACTCCACGAC -3’. QRTPCR was performed using LightCycler 480 SYBR Green□I Master (Roche, 4707516001). Data were collected using QuantStudio 12□K Flex software. mRNA expression was normalized by the expression of either NeoR or GAPDH. Amplification steps were as follow: 95°C for 10’’, 60°C for 10’’ and 72°C for 10’’(40x cycles).

### H&E and Immunofluorescence

Lungs were collected and fixed in 4% PFA overnight and processed for paraffin-embedding. Slides were stained for Haematoxylin and Eosin (H&E) using standard protocols and tumour burden was quantified as tumour area against total lung area. For immunofluorescence slides were stained with were stained with αCD4 (abcam, Ab23990), αCD8 (abcam, Ab217344), αEGFP (Invitrogen, A6455) and DAPI (D9542, Sigma). Confocal microscopy pictures were taken with a Leica STELLARIS microscope.

### Whole genome sequencing and RNAseq analysis

All downstream analyses were performed on WGS and RNA-seq data from Mlh1-KO lung tumour in mouse models and their parental control (ABA19933). Reads were quality-checked with FastQC and aligned to mm10 (GRCm38) with BWA-MEM v0.7.17 (default parameters, 16 threads). Per-sample BAMs were merged (samtools merge), duplicates marked (samtools markdup), and sorted/indexed (samtools sort/index, v1.13). Somatic SNVs and indels were called in parallel via Strelka2 v2.9.10, using the 30 days after the knockout sample (ABA19933) KO-30 BAM as “normal” and filtering out multi-nucleotide variants.

Per-sample VCFs were merged and subsetted to include only those variants that passed Strelka filters (FILTER=PASS). In addition to observed SNVs, we also simulated all three possible base substitutions per site. Both simulated and observed variants were annotated with ANNOVAR (2020Jun07) using the refGene database (mm10).

To quantify immune-specific selection pressure across mutational contexts, we implemented a Bayesian framework for modeling immune-related dN/dS ratios. Mutation data were annotated with trinucleotide context and mutation type. Functional classification was performed using ANNOVAR, while trinucleotide context was derived from genomic coordinates and the mm10 (GRCm38)) reference genome. Mutations were subsequently partitioned based on MHC presentation of the wild-type peptide that spans the mutated residue. Peptides predicted to bind MHC defined the ‘putatively immunogenic’ group, while others formed the non-presentable group. Binding predictions were generated with NetMHCpan4.1, using its two rank-thresholds to call a peptide a binder.

For each somatic variant, we constructed all possible mutant peptides/amino acid k-mers of length 8 to 12 amino acids overlapping the altered residue. Each amino acid k-mer was scored for predicted binding affinity to the H-2-kB and H-2-Db alleles. A mutation was designated as ‘putatively immunogenic’t if at least one k-mer containing the altered amino acid was predicted to fall within the top 0.5% (or, in an alternative analysis, 1%) of high-affinity binders by NetMHCpan, with all remaining mutations being considered not presentable by MHC.

For each sample, we aggregated synonymous and non-synonymous mutation counts stratified by mutational context (192-trinucleotide context, deriving from the standard 96-channel trinucleotide mutational spectrum, considered on both DNA strands). This was aggregated alongside the corresponding background mutational opportunity (in base-pairs, bps), defined as the number of loci at risk of mutation of that class (synonymous or non-synonymous). To ensure robust estimation in sparse settings, we used the arm package in R to fit a Bayesian generalized linear model (BayesGLM) with a Poisson likelihood and log-link function. The primary formula modeled mutations as “MutationCount ∼ isNS * Target * Context”. Here, isNS (non-synonymous indicator), Target (MHC-binding or non-binding region) and Context were modeled as interacting categorical variables with an offset for log(bps) to normalize for mutation opportunity. Expected mutation opportunities (bps) were calculated once over the whole coding exome and applied to both MHC-binding and non-binding windows to ensure a shared mutational risk background.

Furthermore, for every combination of antigen-presentation threshold, gene-expression filter and treatment group, we took the median of dN/dS estimate values across tumours in that slice. The reported dN/dS range for a given group represents the smallest and the largest of these slice-specific medians, reflecting how much the apparent strength of selection changes across analytic settings. To quantify the treatment effect, the median dN/dS estimates of every treatment slice was divided by the median dN/dS of the matched control (same filter and threshold), yielding a rate-ratio. The minimum and maximum of these ratios constitute the rate-ratio range, indicating how consistently the treatment tightens (ratio < 1) or relaxes (ratio > 1) purifying selection relative to baseline (no treatment).

SUPPA2^41^ was used to calculate the PSI (percent spliced-in) values for four event classes: retained intron (RI), skipped exon (SE), alternative 3’ splice site (A3) and alternative 5’ splice site (A5), where PSI equals the fraction of transcripts including the segment of interest (intron for RI, exon for SE, proximal-site extension for A3/A5). For each event we computed ΔPSI as the difference in mean PSI between treatment and control groups. To test whether the same events behave differently under the two treatments, we compared the ΔPSI distributions for NMDi vs Vehicle and aPD1 versus IgG using a Mann-Whitney U test.

### Construction of training peptide sets and IpLM fine-tuning

Mutational data from NMDi-treated samples were processed to obtain full protein sequences, which were segmented into overlapping 8–12 amino-acid peptides (Figure 5 C). Each peptide was scored for predicted MHC class I binding affinity using NetMHCpan-4.1 for H-2-Kb and H-2-Db alleles. High-affinity binders (NetMHCpan %rank ≤ 1) were further annotated according to the underlying mutation: peptides derived from nonsynonymous mutations were treated as immunogenic-like positives, whereas synonymous mutations yielding identical self-peptides formed the matched negative set. Weaker-rank peptides (sampled from %rank bins 39, 41, 49 and 51) were used for control experiments with non-binders (Figure 5 D); in these control datasets, the classification based on immune reactivity is presumably not feasible.

Our IpLM-Amplify (Immuno-peptide Language Model) models were generated by fine-tuning foundation models from the Amplify architecture^47^. Neural net model training used a standard five-fold cross-validation scheme. IpLM-Amplify models were trained using a standardized pipeline built on the AMPLIFY_120M encoder and a lightweight MLP classification head. Peptides were tokenized using the corresponding AMPLIFY tokenizer and encoded jointly with the backbone. Fine-tuning employed the AdamW optimizer with a base learning rate of 3×10□□, and a batch size of 32. To stabilize optimization, the encoder was kept frozen for the first two epochs before unfreezing all layers for joint training. Models were trained for up to 20 epochs with a ReduceLROnPlateau scheduler monitoring validation loss and with early stopping criteria based on validation loss and PR-AUC. For each cross-validation fold, the best-performing checkpoint was saved and used for downstream evaluation.

For an additional test, the ESM-C 300M protein language model was fine-tuned, thus generating the IpLM-ESMC variety. Here, peptides were encoded with the ESMCClassifier wrapper around the ESMplusplus_small backbone (Synthyra/ESMplusplus_small), using its native tokenizer and a built-in classification head. Data splitting, loss functions, regularization, and evaluation metrics followed exactly the same setup as described above for IpLM-Amplify, including group-aware folds defined at the transcript (NM_ID) level to avoid leakage across variants. The model was optimized end-to-end with AdamW (learning rate 3×10□□, batch size 32, hidden-state dropout 0.2) without staged encoder freezing; a single learning rate was applied to all trainable parameters. Training ran for up to 20 epochs with a ReduceLROnPlateau scheduler on validation loss and the same early-stopping scheme (primary criterion: validation loss; secondary tie-breaker: PR-AUC). For each cross-validation fold, the best-performing checkpoint was retained for subsequent analyses.

The modes were fine-tuned on different peptide sets: (i) observed nonsynonymous (i.e. mutated) vs synonymous (i.e. wild-type) MHC binder peptides, (ii) weaker-rank NetMHCpan i.e. non-binders, for both mutated and wild-type, and (iii) neoantigens simulated using SBS96 mutational signatures, while respecting transcriptional strand asymmetry (i.e. equivalent to SBS192). Model outputs were evaluated against held-out folds using area under the receiver-operating characteristic curve ROC-AUC. Statistical differences in AUC were assessed via fold-wise DeLong tests combined using a weighted Stouffer method to obtain aggregate p-values. To assess external generalization, the final cross-validated IpLM-Amplify models were additionally evaluated on the TESLA consortium neoantigen benchmark dataset^46^. For each fold, predictions were generated using the corresponding held-out model checkpoint, and ROC-AUC was computed to quantify performance on this independent human dataset. Frozen and fine-tuned IpLM-Amplify out-of-fold embeddings were reduced to 50 dimensions using PCA and further projected into two dimensions using UMAP (n_neighbors=15, min_dist=0.1). UMAP embeddings were generated independently for frozen and tuned representations. For each peptide class (positive vs negative), a 2D kernel density estimate (Gaussian KDE) was evaluated on a shared grid for the frozen and tuned embeddings. Density-difference maps were computed as Δ = KDE_tuned − KDE_frozen, using a shared color scale across classes to enable direct comparison. Positives and negatives were analyzed separately to isolate class-specific shifts in representation space induced by fine-tuning.

### Position-wise hydrophobicity enrichment analysis

To characterise positional biochemical differences between immunogenic-like and non-immunogenic-like peptides, both within the training dataset of mutations, and also within the trained neural net of the model IpLM-Amplify, we performed a position-wise residue enrichment analysis on predicted MHC class I ligands of lengths 8–12 amino acids. For the IpLM-Amplify analysis, model-derived labels were obtained by collecting out-of-fold predictions across the five validation folds, ensuring that each peptide was evaluated by a classifier that had not seen it during training.

Peptides were aligned by position (P1–PΩ, where PΩ denotes the C-terminal position of the peptides), and for each position we computed, separately for immunogenic-like and non-immunogenic-like peptides, the proportion of residues belonging to one of three classes: aromatic hydrophobic (F/Y/W), aliphatic hydrophobic (L/I/V/M) and, as a control, charged (D/E/H/K/R). Statistical association between immunogenicity and presence of each residue class at each position was tested using a Cochran–Mantel–Haenszel (CMH) test stratified by peptide length (8–12 aa), ensuring that enrichment estimates were not confounded by differences in length distributions. When only a single peptide length contributed data for a position, the test reduced to a standard Fisher’s exact test. This analysis returned a length-adjusted odds ratio (OR) and p-value for each position and residue class. These statistics were visualized as “spaghetti plots,” (Figure 5 J-K, Supplementary Figure 7 A-D) where immunogenic-like and non-immunogenic-like frequency traces were overlaid along the peptide axis (x axis). Statistically significant positions in difference between immunogenic-like and non-immunogenic like (CMH test p < 0.05) were annotated with significance stars.

### Preprocessing of scRNA-seq and TCR Data

Eight multiomic single-cell samples (four treated and four untreated) were generated using the 10x Genomics Chromium Single Cell 5′ and V(D)J v3 (R2-only) chemistry, enabling simultaneous capture of gene expression and T cell receptor (TCR) sequences from individual cells. TCR libraries were processed using Cell Ranger v8.0.1 (10x Genomics) with the cellranger vdj pipeline and aligned to the mouse reference vdj_GRCm38_alts_ensembl- 7.0.0. Corresponding gene expression libraries were processed with the cellranger count pipeline (v8.0.1), using the GRCm39-2024-A mouse reference transcriptome. Default parameters were used unless otherwise specified.

Quality Control: To ensure high-quality single-cell transcriptomic data, cells were filtered based on standard thresholds. Cells with fewer than 10,000 total UMI counts or fewer than 200 detected unique genes were excluded from downstream analysis. Additionally, cells in which mitochondrial transcripts accounted for more than 5% of total expression were removed, as a high proportion of mitochondrial reads is indicative of low-quality or dying cells. Cells not meeting these criteria were excluded. Putative doublets were identified using the DoubletDetection tool (v4.2)^42^ and removed.

Dimensionality reduction and clustering were performed using Scanpy v1.9.3^43^. Log-normalized and scaled expression data were used to compute principal components (PCA), and batch correction was performed using Harmony^44^, via the harmonypy Python implementation. The top 30 Harmony-adjusted principal components were used for Leiden clustering (resolution = 0.15). Differentially expressed genes per cluster were identified using the Wilcoxon rank-sum test, and clusters were annotated based on known marker gene signatures corresponding to major immune cell types.

The following cell populations were identified: T cells, cycling/proliferating cells, classical monocytes, tissue-resident macrophages, conventional dendritic cells type 2 (cDC2), B cells (including early and mature subsets), natural killer (NK) cells, inflammatory neutrophils, stromal/mesenchymal cells, plasmacytoid dendritic cells (pDCs), and M2-like monocyte-derived dendritic cells (moDCs).

TCR quality control was conducted using the Scirpy package. TCR chain types were indexed using ir.pp.index_chains(mdata), and chain-level metrics were calculated with ir.tl.chain_qc(mdata). Cells containing orphan TCR chains (only α or β) were excluded from subsequent analyses.

Latent dimensionality reduction was performed using scVI (v1.1.6.post2). Highly variable genes were selected within each preliminary cell-type compartment using Scanpy’s highly_variable_genes function (parameters: min_mean=0.0125, max_mean=3, min_disp=0.5), resulting in 15,150 input genes. The scVI model was trained on raw counts, using sample identity as a batch key. The model architecture included two encoder/decoder layers (n_layers=2) and a 30-dimensional latent space (n_latents=30). Training was performed with early stopping and terminated when the ELBO validation loss plateaued (45 epochs). Leiden clustering was applied to the latent space (resolution = 1.0), and marker genes for each cluster were identified using the Wilcoxon rank-sum test. Final cell type annotation was assigned based on cluster-specific gene expression profiles.

To assess the functional states of CD8□ T cells, gene signature scores were calculated using the score_genes function in Scanpy. Specific gene sets were used to define distinct CD8□ T cell programs: an exhaustion signature (Pdcd1, Lag3, Havcr2, Klrg1, Tigit, Cd160, Btla, Ctla4, Entpd1, Id2), an effector signature (Ifng, Tnf, Gzmb, Cxcl9, Cxcl10, Ccl3, Ccl4, Tbx21, Prf1), and a cytotoxic signature (Prf1, Gzma, Gzmb, Gzmk, Nkg7, Klrc1, Klrd1, Klrk1, Il2rb). These scores enabled comparative profiling of CD8□ T cell functional states across experimental.

### Integrative genomic, transcriptomic and functional analysis for prioritization of NMD targets

Matched tumour-normal Whole-Exome Sequencing (WES), RNA-sequencing (RNA-seq), and gene-level Copy Number Alteration (CNA) data were obtained from The Cancer Genome Atlas (TCGA)^15^ via the Genomic Data Commons (GDC) portal (https://portal.gdc.cancer.gov/). Sequencing data processing was performed as previously described^19^. Tumour NMD efficiency (NMDeff) scores were retrieved from, utilizing two orthogonal metrics: the “ETG” method (based on mRNA expression of endogenous NMD targets versus non-NMD controls) and the “ASE” method (based on allele-specific read counts of germline nonsense and frameshift mutations in NMD-triggering regions).

To assess the functional impact of NMD factor alterations, we utilized generalized linear regression models associating somatic events in 15 NMD genes (core: UPF1, UPF2, UPF3A, UPF3B, SMG1, SMG5–9; EJC: EIF4A3, CASC3, MAGOH, MAGOHB, RBM8A) with NMDeff scores. Analyses were performed at the pan-cancer level and within 31 individual cancer types; cancer types with mean tumor purity < 0.75 were excluded. Somatic alterations were stratified into three categories: (i) Truncating/Missense mutations, defined as truncating (stopgain, frameshift insertion or deletion, splicing) or missense (non-synonymous, start/stop-loss, non-frameshift insertion or deletion) variants, requiring >= 2 observed variants across samples; (ii) CNA gain (GISTIC2 score > 0); and (iii) CNA deletion (GISTIC2 score < 0). Samples harboring somatic SNVs in the queried gene were excluded from CNA analyses to isolate copy-number effects.

Multivariate models were adjusted for the following covariates: tumor purity, tumor mutation burden (TMB), cancer subtype (NMF mRNA-seq clustering), CNA burden, patient sex and age, RNA-seq library size, and 86 principal components representing arm-level CNA changes^19^. Covariates were dynamically excluded where applicable (e.g., sex in single-sex cancers or variables with excessive NAs). For ranking, genes were ordered by association p-value within each cancer type and somatic alteration category. Associations yielding negative beta coefficients (indicating reduced NMD efficiency) were prioritized. A final global priority rank was derived by averaging the gene rankings across all 30 cancer types and the pan-cancer cohort.

We correlated gene expression levels (log2(TPM+1)) of the 15 NMD genes with NMDeff scores at the pan-cancer level, performing separate analyses for ASE and ETG metrics. Prior to correlation, expression data were adjusted for potential confounders related to tumour microenvironment composition using the removeBatchEffect function from the limma R package. The correction model included tumour purity and the fractions of 9 common cell types as covariates (hematopoietic cells, leukocytes, lymphocytes, B-cells, T-cells, macrophages, endo-epithelial cells, endothelial cells, fibroblasts), with cell proportions derived using the UniCell: Deconvolve Base (UCDBase) bulk RNA-seq deconvolution method^52^. To ensure that tissue-specific expression patterns were not removed during correction, cancer type was included in the design matrix to preserve biological heterogeneity across cohorts. Finally, Pearson coefficients of determination (R2) were normalized to a [0–1] range using min-max scaling: (R2 - R2min) / (R2max - R2min).

To evaluate functional co-dependencies between NMD factors, we leveraged large-scale genetic interaction data from the Achilles Project, accessed via the DepMap portal^53^. This included gene-level siRNA screens (712 cell lines × 17,310 genes) and CRISPR-Cas9 knockout screens (957 cell lines × 18,018 genes). To enhance analytical robustness and mitigate technical biases, we additionally utilized a corrected dataset processed via the “Onion” normalization pipeline and Robust Principal Component Analysis (RPCO)^26^. From this pipeline, we retrieved the symmetrical Similarity Network Fusion (SNF) score matrix representing 18,119 genes (”snf_run_rpca_7_5_5.Rdata”, https://zenodo.org/records/7671685#.Y_gi9nbMK5c).

For the standard siRNA and CRISPR datasets, we calculated correlations between the fitness profiles of 14 NMD genes and UPF1. For the RPCO-corrected dataset, pre-computed SNF similarity scores were utilized. To facilitate integration across the three heterogeneous datasets, scores were normalized: correlation coefficients underwent Min-Abs scaling, while SNF scores underwent Min-Max scaling. A final composite dependency score was derived by averaging the normalized values across all three datasets, with genes ranked by this mean score (Rank 1 indicating the strongest functional codependency with UPF1).

We performed Pearson correlation analyses between the mRNA expression of 15 NMD genes and CD8□ T-cell proportions for 20 cancer types and pan-cancer. Immune cell fractions were estimated using the CIBERSORT “LM22” signature and retrieved from The Cancer Immunome Atlas (TCIA) (https://tcia.at/home)^54^. Prior to analysis, gene expression data were corrected to account for tumour purity and three non-immune stromal lineages (epithelial cells, endothelial cells, and fibroblasts) to preserve immune-related expression variance for correlation.

### Data and code availability

ScRNA-seq data, WGS and bulk RNAseq data and analysis codes are available upon request. Any additional information required to reanalyse the data reported in this paper is available upon request.

## Supplementary Figure Legends

**Supplementary Figure 1.** A) Distribution of statistical associations (p-values) between somatic alterations and individual NMD efficiency (iNMDeff) for 10 core NMD genes (5 EJC genes were excluded for better visualization) across 31 TCGA cancer types and pan-cancer. Genes are ordered by their median association significance. Points are coloured according to the gene’s relative rank among 15 NMD genes within that specific cohort (colour gradient: 1 = top rank/strongest association to 15 = lowest rank/weakest association). Statistical significance of the difference in median distributions between SMG5, SMG7, UPF1 and SMG1 was determined using a one-tailed Mann–Whitney U test. B) Pan-cancer associations between the mRNA gene expression of 15 NMD factors and iNMDeff. The plots display the Pearson coefficient of determination (R2), Min-Max normalized to a 0–1 scale, for allele-specific expression (ASE, left) and endogenous target gene (ETG, right) NMDeff methods. Asterisks indicate statistical significance of the correlations: *p < 0.05, **p < 0.01, ***p < 0.001. C) Assessment of functional co-dependency between NMD factors and the core helicase UPF1. The plot displays relative functional interaction scores (y-axis), calculated by integrating Pearson correlations (Abs-Max normalized) from Achilles CRISPR and RNAi screens with SNF similarity probabilities (Min-Max normalized) from the RPCO-corrected dataset. Genes are ordered by their mean normalized score across these three platforms. D) Detailed breakdown of functional co-dependency scores stratified by dataset (Achilles CRISPR, Achilles RNAi, and RPCO). Genes within each facet are reordered according to their dataset-specific normalized interaction score, highlighting the strongest functional partners of UPF1 within each individual screening platform. E) Heatmap displaying Pearson correlation coefficients between the expression of 15 NMD genes and CD8+ T-cell proportions (estimated via CIBERSORT) across 20 cancer types and the pan-cancer cohort. SMG7 and SMG1 exhibit the most consistent negative association with CD8+ T-cell infiltration; this aligns with the expectation that elevated expression of these genes drives NMD activity, thereby suppressing immune infiltration.

**Supplementary Figure 2.** A) Tumour response data are derived from individual growth curves shown in Figure 1E. Responders were defined as animals exhibiting a ≥30% reduction in tumour growth relative to controlś average weight. The pie charts illustrate the proportion of responders (coloured segments) and non-responders (grey). Absolute counts are shown. C) Gating strategies used for analysis of GFP ^+^ cancer cells. D) Wet weight (mg) of the lung of vehicle/IgG2a (n=3), αPD1 (n=5), NMDi (n=2), or NMDi+aPD1 (n=3) treated mice. p values were calculated by one-way ANOVA E) Combination therapy elicits a distinct transcriptional program beyond either αPD1 or NMDi inhibition alone. A total of 222 genes whose absolute log2 fold-change exceeded 0.5 in at least one comparison is displayed for αPD1 (n=4), NMDi (n=4), or NMDi+ αPD1 (n=3) compared to the vehicle/IgG2a (n=5), each contrasted to the pooled Vehicle + IgG control. Rows are hierarchically clustered (Pearson correlation). Colour denotes log2FC, with blue indicating downregulation, white no change, and red indicating up-regulation.

**Supplementary Figure 3.** A) Percentage of CD8□ cells in vehicle (n=4) and NMDi (n=4) groups before the injection of αCD8 (Pre-treatment) and at the start of NMDi therapy 24 hours later (treatment). p-value was calculated by two-tailed unpaired *t*-test -B) Gating strategies used for analysis of CD8^+^ T cells.

**Supplementary Figure 4.** A) Quality control metric distribution across samples. (Top left) Total UMI counts per cell. (Top right) Number of unique genes detected per cell. (Bottom left) Percentage of mitochondrial gene counts per cell. (Bottom-right) Percentage of ribosomal gene counts per cell B) Top differentially expressed genes per cell type. Dot plot displaying the top 3 differentially expressed genes (log fold-change > 2.5) per cell type across all major annotations. C) Top differentially expressed genes in major immune compartments. Dot plot showing the top differentially expressed genes (log fold-change > 2) per cell subtype across three compartments: T cells (top), macrophages (middle), and neutrophils (bottom).

**Supplementary Figure 5.** Immune dN/dS across treatment under two expression filters. A) Log-scaled immune dN/dS for Control (IgG2a), αPD1, NMDi, and aPD1+NMDi without any expression filter (all genes included). B) Same analysis restricted to genes with TPM > 1in the corresponding tumour. In each panel every point is one tumour, violins depict the kernel density and embedded box-and-whisker plots show the median. The dashed horizontal line indicates neutral expectation (log dN/dS = 0). Within each panel, the left subplot shows mutations whose peptides are in the top 0.5% binders predicted by NetMHCpan4.1 (%rank = 0.5), and the right subplot shows the broader set with peptides in the top 1% binders (%rank = 1). C) dN/dS-cv analysis identifies treatment-specific positive selection among expressed genes. For five treatment arms (Vehicle, IgG, αPD1, NMDi, αPD1+NMDi) the -log10(q) statistic from dNdScv is shown only for genes with detectable expression (raw TPM > 0). The panel shows 20 most significant expressed genes per arm. Coloured lines trace -log10(q) for each treatment. The red dashed line marks the false-discovery rate threshold (FDR = 0.05, -log10=1.30). Upward triangles highlight genes that surpass this threshold in the corresponding arm, indicating positive selection. Gene labels on the x-axis list the mouse symbol followed by the corresponding human orthologue in parentheses.

**Supplementary Figure 6.** NMDi inhibition and PD-1 blockade drive directional splicing shifts across four classes of alternative splicing events. Volcano plots of ΔPSI versus raw p-values (Welch two-sided t-test) are shown for A) retained introns (RI), B) skipped exon (SE), C) alternative 3’ splice sites (A3) and D) alternative 5’ splice sites (A5). In the left columns each event type compare NMDi-treated versus Vehicle controls while in the right column the same event types compare aPD1 versus IgG control. On each plot the x-axis is ΔPSI (mean treatment - mean control) and the y-axis is -log10(raw p-value). A blue dashed horizontal line marks p=0.05, and green dashed vertical line mark |ΔPSI|=0.1 (10% splicing change). Frame-shifting (FS) events are shown as red circles; in-frame (IF) events as blue squares; all other points are rendered in light grey to indicate non-significant or sub-threshold changes.

**Supplementary Figure 6.** Position-wise enrichment of hydrophobic aliphatic and charged residues in immunogenic-like versus non-immunogenic-like peptides.

**Supplementary Figure 7.** A) Aliphatic hydrophobic residue enrichment analysis using observed mutation labels (“true” labels), comparing peptides generated by nonsynonymous versus synonymous mutations. Red and blue curves show residue frequencies at TCR-facing (non-anchor) positions for immunogenic-like and self-like peptides, respectively, while grey curves indicate anchor and flanking positions. Asterisks denote positions with significant associations between peptide class and aromatic residue presence (*p < 0.05; Cochran–Mantel–Haenszel test stratified by peptide length). B) Same analysis as in A, but for charged amino acid residues. C-D) Same analysis as in A and B, but using IpLM-Amplify classifier–predicted labels for the same NMDi-derived high-affinity binder peptides used in Figure 5 H. E) KDE difference for immunogenic-like peptides showing the change in 2D density of positive peptides after fine-tuning (tuned - frozen). The red indicates regions of increased density and blue indicates decreased density. F) Same KDE difference plot as in E, showing the corresponding density shift for negative peptides.

## Authors’ contributions

I.Z., D.O., D.K. Conceptualization. Data curation. Formal analysis. Investigation. Methodology. Validation. Visualization. Project administration. Writing - original draft. Writing - review & editing. M.A., E. A., L.O., A.B., Y. K., C.M. M., E.M., G.P. Data curation. Formal analysis. Methodology. Visualization. Writing - original draft. E.M. Data curation. Formal analysis. Methodology. Visualization. Writing - original draft. Supervision. FS and A.J. Conceptualization. Visualization. Project administration. Writing - original draft. Writing - review & editing. Supervision. Resources. Funding acquisition.

## Acknowledgements

We thank Dr A. Celià-Terrassa, Dr. K. Sutherland and Dr I. Toledano for providing reagents; CRG/UPF Flow Cytometry Unit with sample preparation and analysis. This work was supported by grants and fellowships from the Spanish Ministry of Science and Innovation Grant to A.J. (PID2021-127710OB-I00, CNS2023-144819), “La Caixa” foundation (51110009 and HR22-00402). A.J. is supported by Ramon y Cajal Research Fellowship (RYC2018-025244-I). I.Z. is funded by an AECC Postdoctoral Fellowship (POSTD234858ZADR). This work was made possible through the “Unidad de Excelencia María de Maeztu’’ funded by the MCIN and AEI (CEX2018-000792-M). Work in the lab of F. S. was supported by an ERC CoG “STRUCTOMATIC” (101088342), Horizon2020 project “DECIDER” (965193), Horizon EU project “LUCIA” (101096473), ICREA professorship, CaixaResearch project “POTENT-IMMUNO” (HR22-00402), Novo Nordisk Fonden Start Package grant, and a Danish Cancer Society project grant.

