## Supplementary material for "Dual inhibition of the nonsense mediated mRNA decay enhances tumour immunogenicity, drives immunoediting, and potentiates checkpoint blockade": Supp Figures

#### Supplementary Figure 1

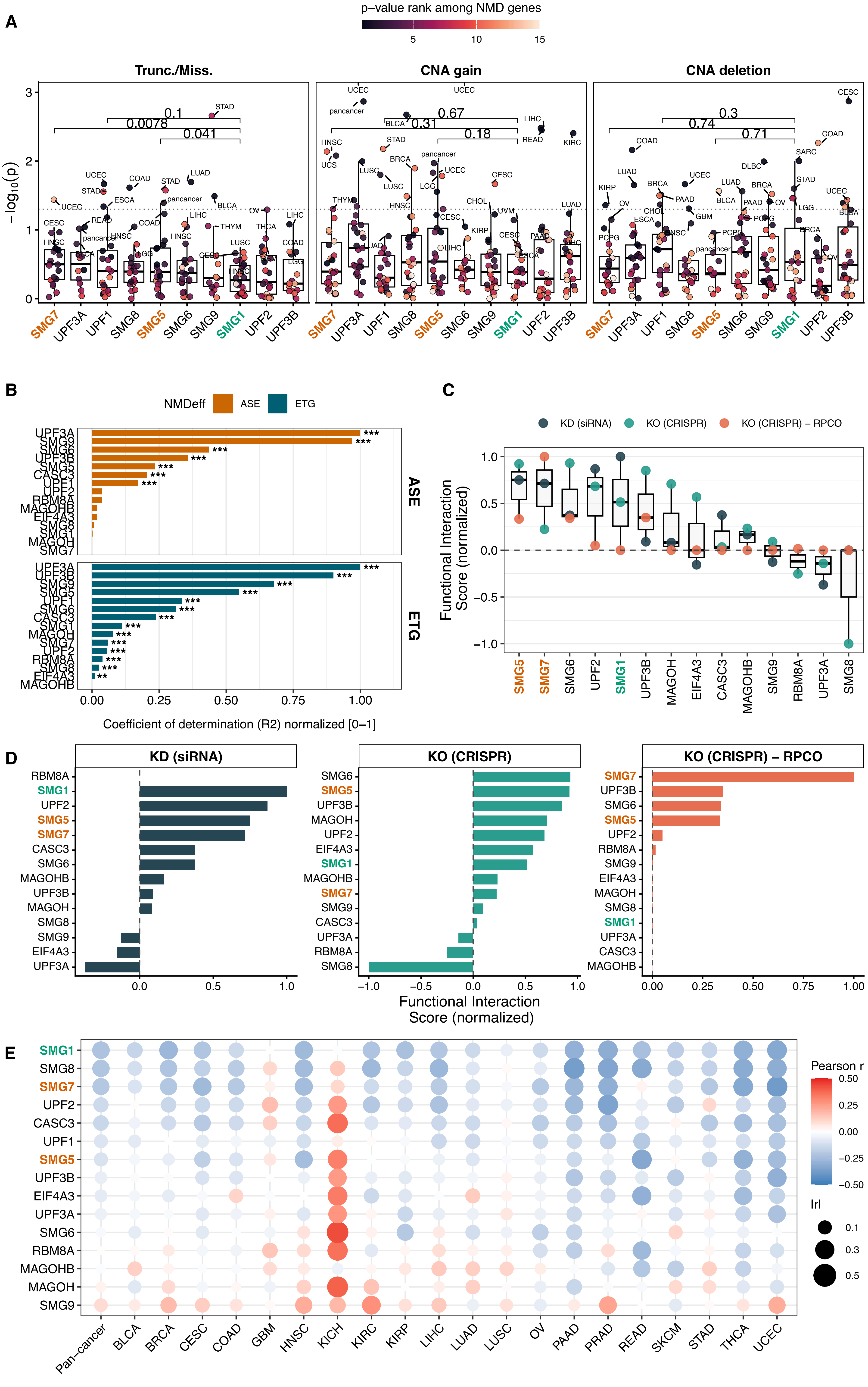

### Supplementary Figure 2

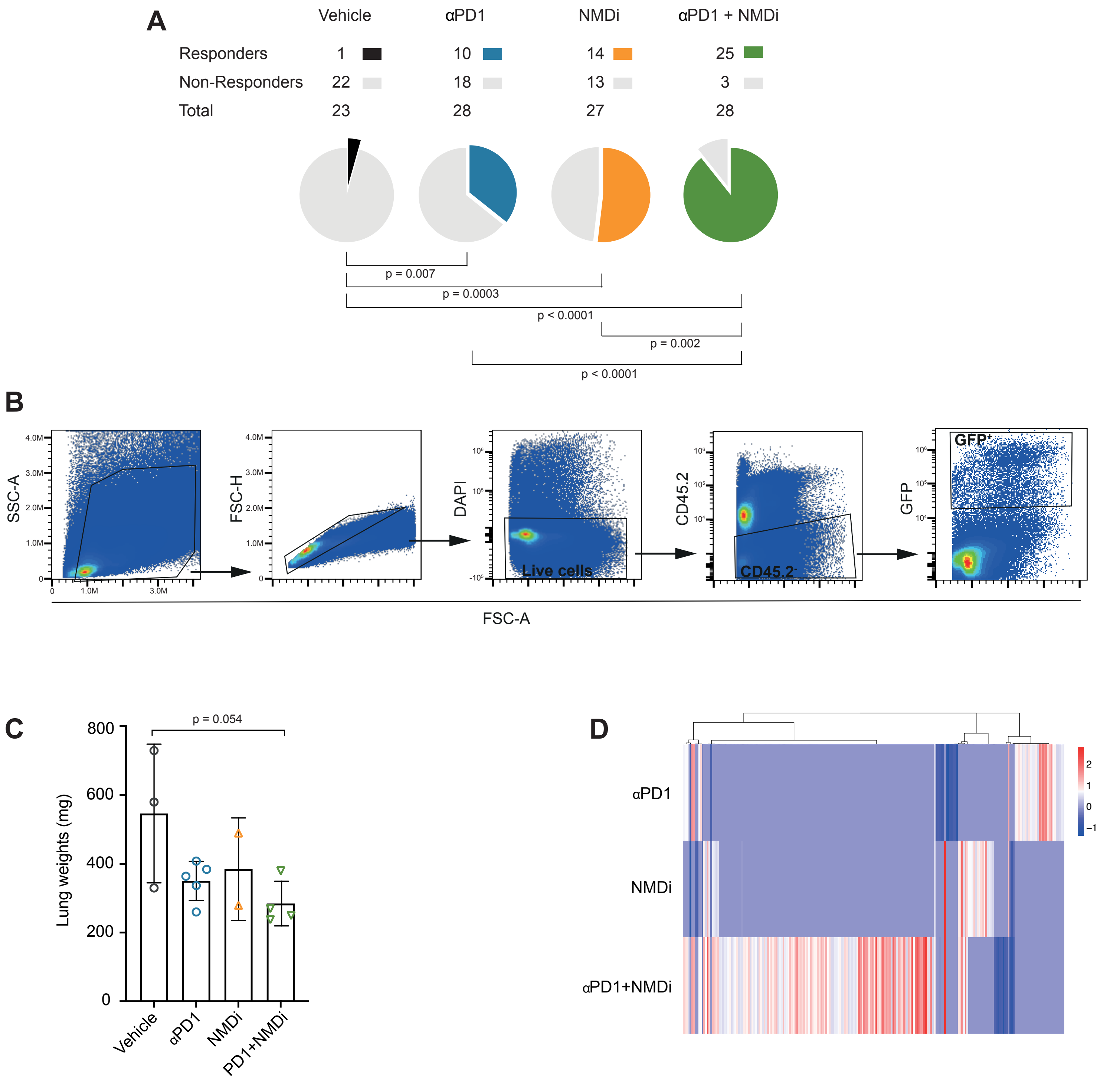

### Supplementary Figure 3

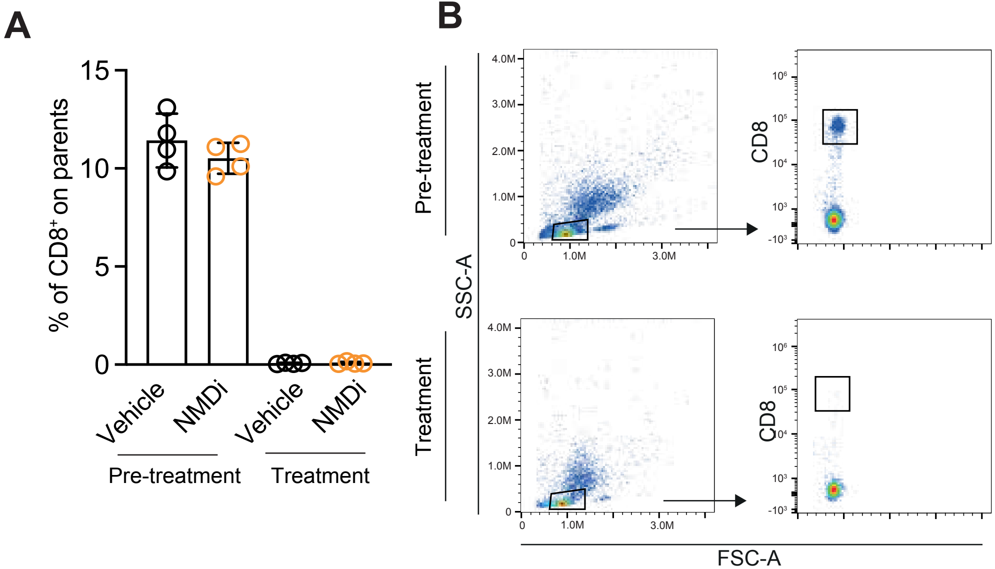

### Supplementary Figure 4

A

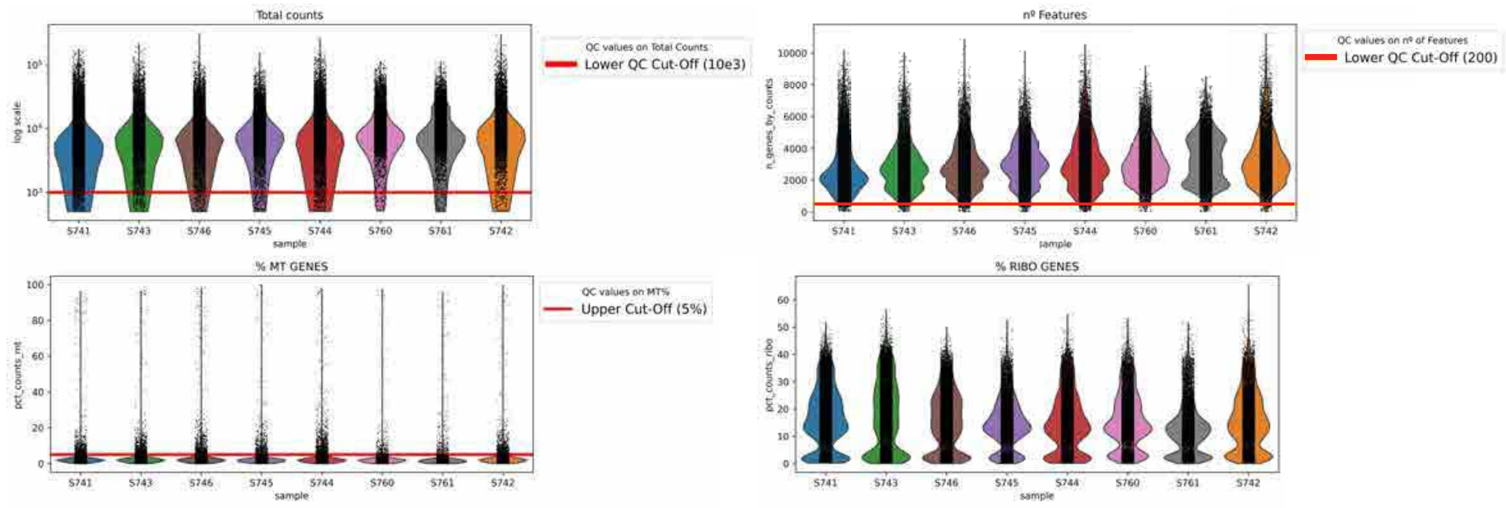


B

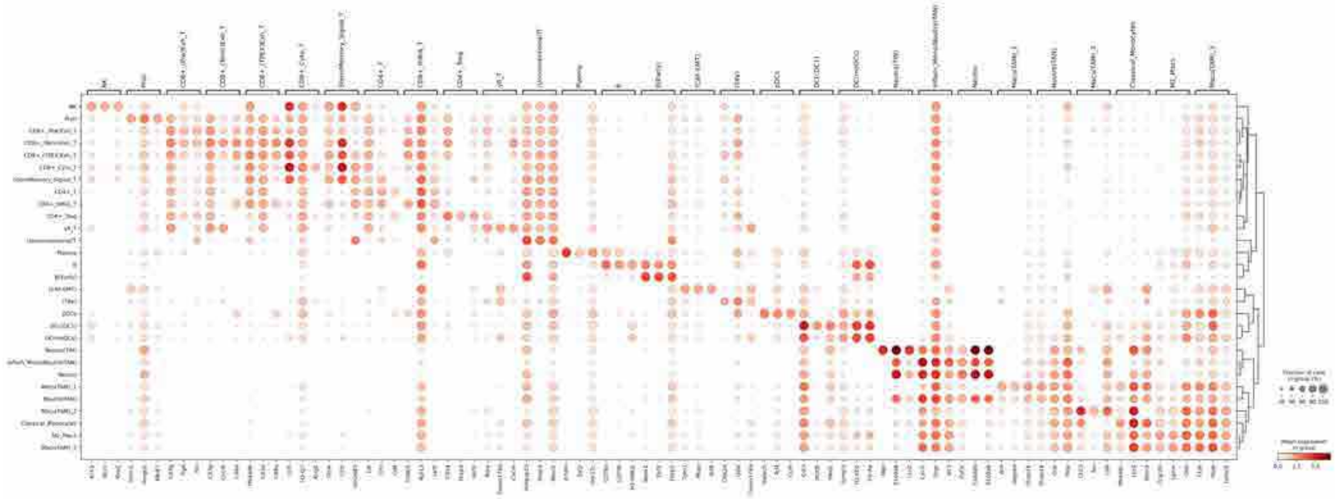

C

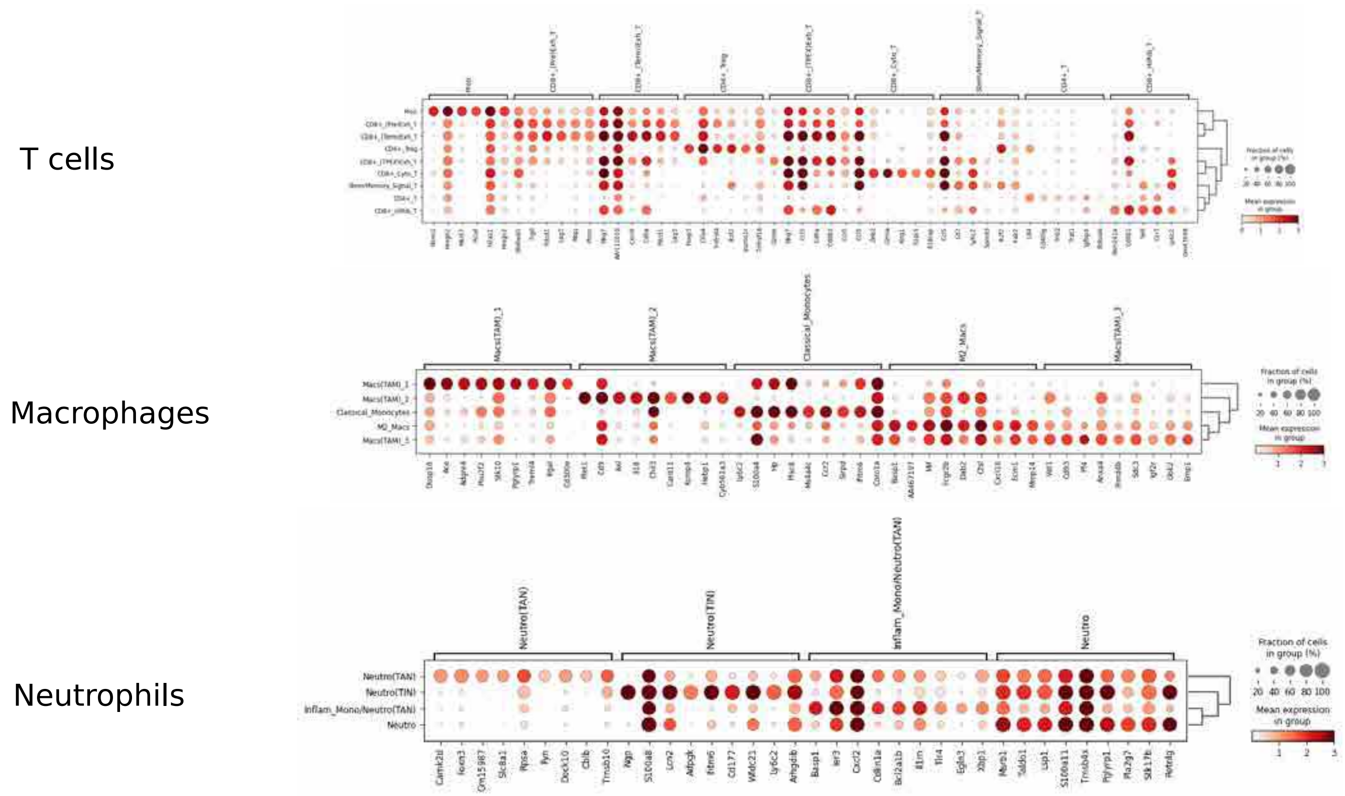

### Supplementary Figure 5

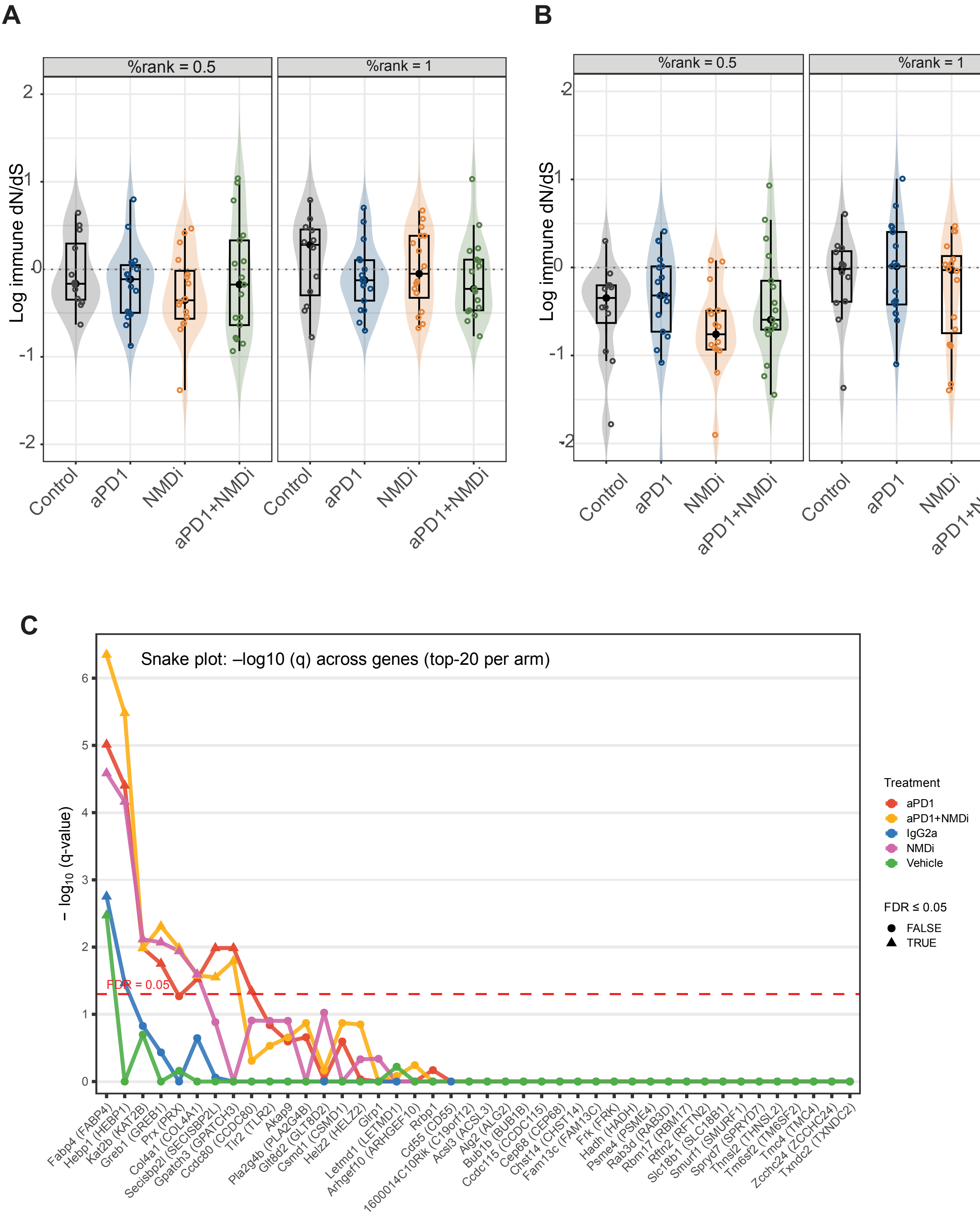

### Supplementary Figure 6

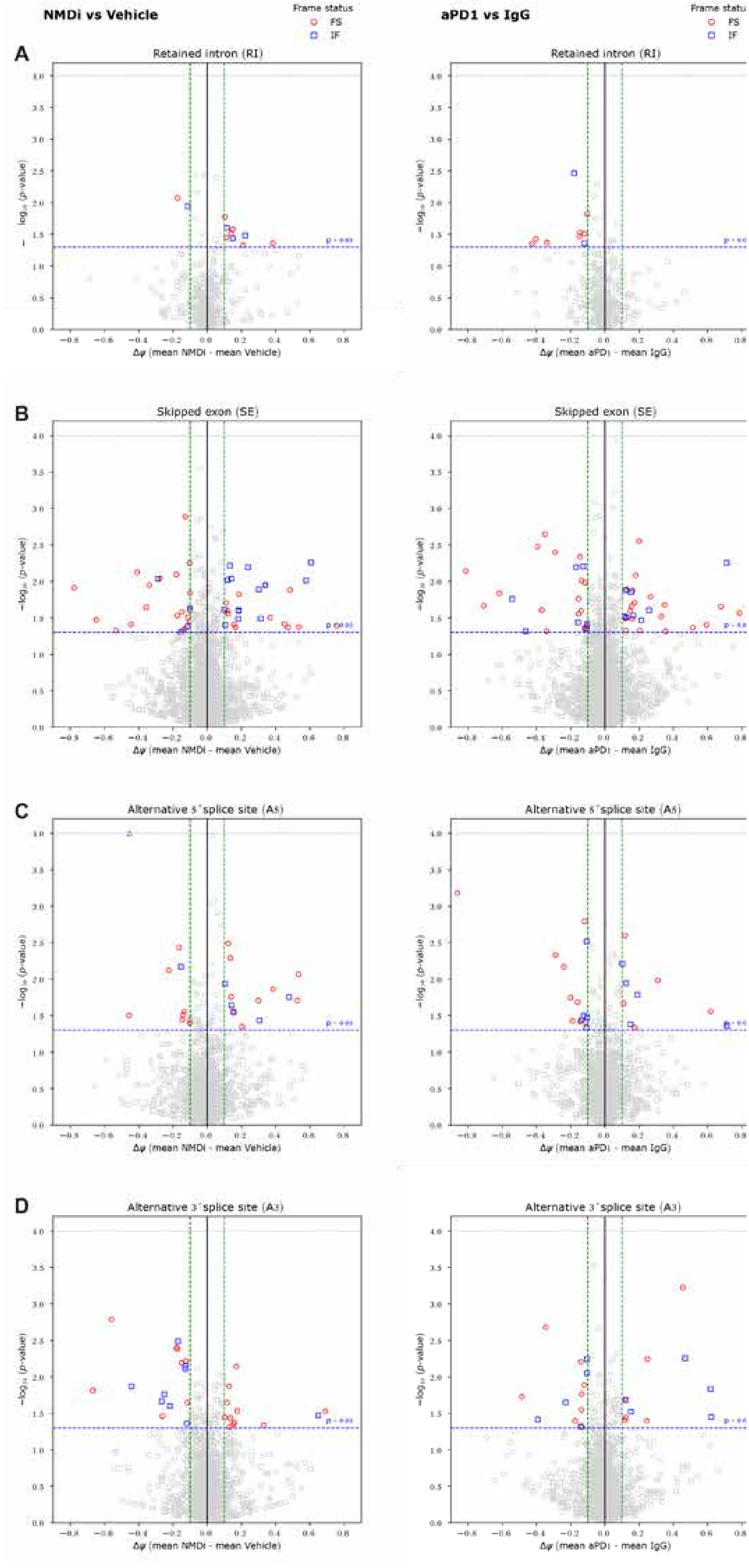

### Supplementary Figure 7

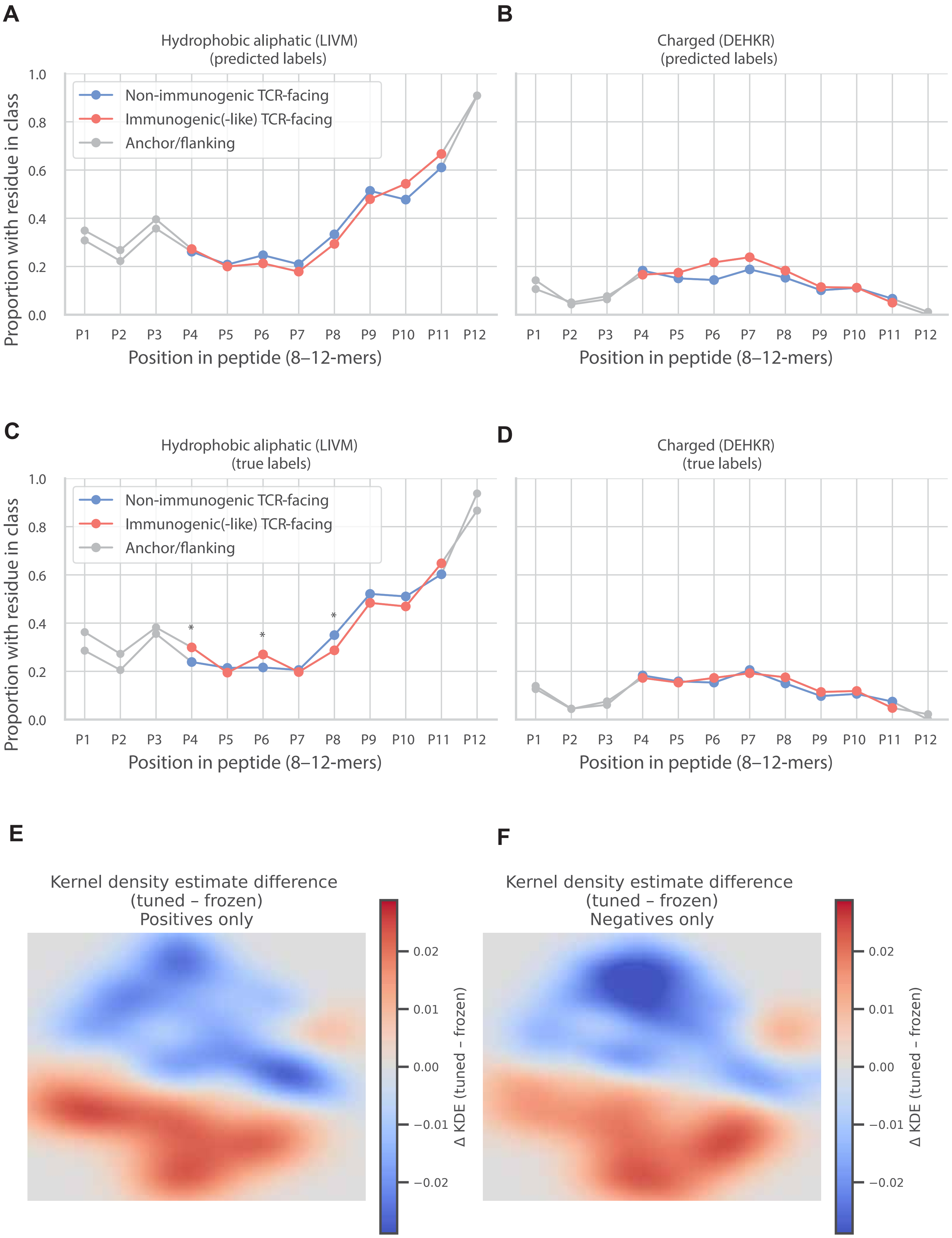
